# The chromatin remodeler ISWI acts during *Drosophila* development to regulate adult sleep

**DOI:** 10.1101/2020.08.11.247080

**Authors:** Naihua N. Gong, Leela Chakravarti Dilley, Charlette E. Williams, Emilia H. Moscato, Milan Szuperak, Qin Wang, Matthew Jensen, Santhosh Girirajan, Tiong Yang Tan, Matthew A. Deardorff, Dong Li, Yuanquan Song, Matthew S. Kayser

**Affiliations:** Department of Psychiatry, Perelman School of Medicine at the University of Pennsylvania, Philadelphia, PA 19104, USA; Raymond G. Perelman Center for Cellular and Molecular Therapeutics, Children’s Hospital of Philadelphia, Philadelphia, PA 19104, USA; Department of Biochemistry and Molecular Biology, Pennsylvania State University, University Park, PA 16802, USA; Bioinformatics and Genomics Program, Huck Institutes of the Life Sciences, Pennsylvania State University, University Park, PA 16802, USA; Department of Anthropology, Pennsylvania State University, University Park, PA 16802, USA; Victorian Clinical Genetics Services, Murdoch Children’s Research Institute, Melbourne, Australia; Department of Paediatrics, University of Melbourne, Melbourne, Australia; Division of Human Genetics, The Children’s Hospital of Philadelphia, Philadelphia, PA 19104, USA; Department of Pediatrics, University of Pennsylvania Perelman School of Medicine, Philadelphia, PA 19104, USA; Department of Pathology and Laboratory Medicine, Children’s Hospital of Los Angeles, Los Angeles, CA 90027, USA; Center for Applied Genomics, Children’s Hospital of Philadelphia, Philadelphia, PA 19104, USA; Department of Pathology and Laboratory Medicine, University of Pennsylvania, Philadelphia, PA 19104, USA; Department of Neuroscience, Perelman School of Medicine at the University of Pennsylvania, Philadelphia, PA 19104 USA; Chronobiology and Sleep Institute, Perelman School of Medicine at the University of Pennsylvania, Philadelphia, PA 19104 USA

**Author notes:** These authors contributed equally to this work (Naihua N. Gong, Leela Chakravarti Dilley).

## Abstract

Sleep disruptions are among the most commonly-reported symptoms across neurodevelopmental disorders (NDDs), but mechanisms linking brain development to normal sleep are largely unknown. From a *Drosophila* screen of human NDD-associated risk genes, we identified the chromatin remodeler *Imitation SWItch/SNF* (*ISWI*) to be required for adult fly sleep. Loss of *ISWI* also results in disrupted circadian rhythms, memory, and social behavior, but *ISWI* acts in different cells and during distinct developmental times to affect each of these adult behaviors. Specifically, *ISWI* expression in type I neuroblasts is required for both adult sleep and formation of a learning-associated brain region. Expression in flies of the human *ISWI* homologs *SMARCA1* and *SMARCA5* differentially rescue adult phenotypes. We propose that sleep deficits are a primary phenotype of early developmental origin in NDDs, and point towards chromatin remodeling machinery as critical for sleep circuit formation.

## Introduction

Neurodevelopmental disorders (NDDs) are highly prevalent and diverse diseases related to abnormal brain maturation. While numerous behavioral phenotypes are commonly associated with individual genetic mutations in NDDs(*1, 2*), sleep disturbances are pervasive across NDDs, and are a significant stressor for individuals and caretakers alike(*3, 4*). Strong clinical associations between disrupted sleep and other NDD symptoms(*5, 6*) suggest that sleep disturbances may be secondary to broader cognitive or behavioral deficits(*7–9*), and are therefore refractory to treatment. Alternatively, sleep dysfunction in NDDs might represent a core phenotype directly related to pathological developmental processes(*10*). As sleep is important for normal neurodevelopment and function(*11*), early sleep disturbances might exacerbate other behavioral issues. Given the high prevalence and significant burden of NDD-associated sleep problems, understanding the mechanistic underpinnings of sleep disruptions is crucial for developing therapeutic interventions.

Sleep in the genetically tractable model organism, *Drosophila melanogaster*, has the defining behavioral characteristics of vertebrate sleep and is regulated by evolutionarily conserved signaling pathways(*12*). These characteristics position *Drosophila* as an ideal, high-throughput model to 1) identify causative NDD risk genes that affect sleep and 2) investigate how these same risk genes may contribute to behavioral pleiotropy. To identify mechanisms underlying NDD-associated sleep disturbances, we screened for sleep abnormalities using RNAi targeting *Drosophila* homologs of human NDD risk genes. Constitutive knockdown of *Imitation SWItch/SNF* (*ISWI*) led to dramatic sleep disturbances in the adult fly. Across species, ISWI and its homologs are ATP-dependent chromatin remodelers that regulate the expression of genes important for neural stem cell proliferation and differentiation(*13–18*). Rare variants in the human homologs of *ISWI*, *SMARCA1* and *SMARCA5* (unpublished data), have been implicated in several NDDs(*18–21*). Moreover, large-scale genome wide and exome sequencing studies on patient cohorts have shown that genetic factors contributing to NDDs converge on chromatin regulation pathways(*22, 23*). Chromatin dynamics are critical for appropriate gene expression during key developmental timepoints(*24*). Thus, dysfunction of these important gene regulatory hubs likely results in a multitude of downstream biological effects, contributing to behavioral pleiotropy seen in NDDs. Delineating how chromatin remodelers like *ISWI* control development of neural circuits involved in diverse behaviors will deepen our understanding of behavioral pleiotropy in NDDs.

In addition to sleep deficits, we found that knockdown of *ISWI* leads to circadian abnormalities in the adult fly, as well as memory and social dysfunction. Temporal mapping revealed *ISWI* acts during dissociable pre-adult stages and spatially distinct circuits to affect these different adult behaviors. At the circuit level, *ISWI* knockdown disrupts the morphology and function of the adult sleep-regulatory dorsal fan-shaped body (dFB) neurons, likely by affecting the cell fate of dFB neurons. Expressing either human *SMARCA1* or *SMARCA5* in the setting of ISWI knockdown differentially rescued adult deficits; specifically, *SMARCA5* but not *SMARCA1* was able to rescue adult fly sleep in the setting of *ISWI* knockdown. Our results delineate how mutations in a single NDD risk gene give rise to primary disruptions of sleep circuit development in the setting of behavioral pleiotropy.

## Results

### ISWI is necessary for normal sleep in Drosophila

In order to identify NDD risk genes with strong effects on sleep, we took advantage of high-throughput sleep assays in *Drosophila*(*25, 26*). We focused on human genes within loci of interest that have been strongly associated with risk for NDD(*1, 27–29*). These loci included chromosomal copy number variants (CNVs) as well as individual risk genes. We performed a reverse genetic RNAi-based screen of *Drosophila* orthologs of NDD-associated human genes (**Fig 1A**) using the *elav-GAL4* enhancer to drive expression of *UAS-RNAi* lines in the developing and adult nervous systems. We individually knocked down 218 genes, comprising a total of 421 unique RNAi lines (including 73 lethal lines) (**Fig 1B-D**). From this screen, we found knockdown of *Imitation Switch/SNF* (*ISWI*) dramatically decreased total sleep duration (**Fig 1E,F**), with the strongest effect during the night (**Fig 1G**). Pan-neuronal *ISWI* knockdown also led to more fragmented sleep, due primarily to a reduction in sleep bout length during the day and the night (**Fig 1H,I**). Although knockdown of several other genes also resulted in increased sleep fragmentation (**Fig 1C, D**), we chose to focus on *ISWI* given its additional involvement in total sleep duration. Knockdown with an independent RNAi line for *ISWI* recapitulated the observed sleep deficits (**Fig 1E,F; Fig S1A-C**). We validated that both tested RNAi constructs decreased *ISWI* mRNA levels (**Fig S2A**), and found that co-expression of a FLAG- and HA-tagged RNAi-resistant *UAS-ISWI* (*UAS-ISWI^Res^-FH*) in the setting of *ISWI* knockdown rescued sleep deficits (**Fig S2B-F**). These results demonstrate that sleep deficits are specific to the effects of *ISWI* RNAi-based knockdown. Sleep homeostasis was also impaired in *ISWI* RNAi flies: *elav-GAL4 > UAS-ISWI RNAi* flies exhibited ∼300 minutes of sleep loss in response to overnight mechanical deprivation (**Fig S1D,E**), but in contrast to genetic controls, failed to exhibit sleep rebound (**Fig 1J; Fig S1D,F**). Thus, *ISWI* knockdown results in decreased and fragmented sleep, as well as deficits in homeostatic rebound.

**Fig. 1:**
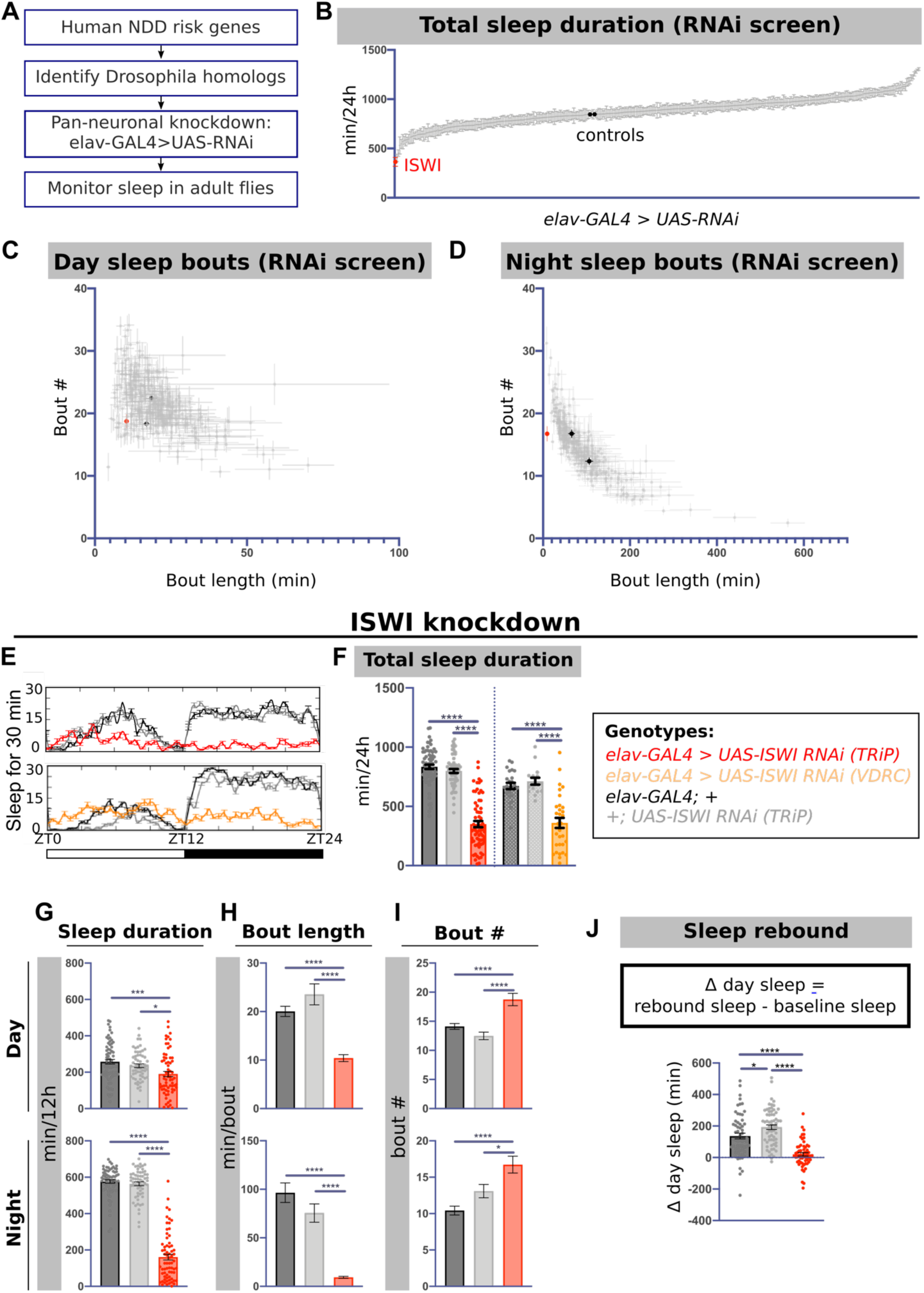
*ISWI* knockdown results in sleep deficits in adult *Drosophila.* A) Design of NDD risk gene sleep screen. B) Total sleep in all viable screened RNAi lines (348 lines, n ≥ 16 per RNAi line). Average sleep bout length and sleep bout number plotted for all lines across the C) day and D) night. E) Representative sleep traces of genetic controls (black, gray) and pan-neuronal *ISWI* knockdown with two independent RNAi lines (red, orange). F) Total sleep, G) day and night sleep, H) average sleep bout length and I) number during the day or night in *ISWI* knockdown compared to genetic controls (n = 70, 61, 68, 24, 16, 31 for groups from left to right in F). J) Comparison of differences in rebound and baseline day sleep in *ISWI* knockdown (red) and genetic controls (gray) (n = 63, 65, 61 from left to right). For graphs in this figure and all other graphs unless otherwise stated: data are presented as mean ± SEM. \**P* < 0.01, \*\**P* < 0.01*, ***P* < 0.001*, ****P* < 0.0001 and analyzed with one-way ANOVA with post-hoc Tukey’s multiple comparison test (B,F-J).

### ISWI knockdown impairs circadian rhythmicity, memory, and courtship behaviors

Patients with NDDs exhibit myriad behavioral disruptions in addition to sleep problems, such as circadian disturbances(*30*), intellectual disability (ID)(*31*), and social deficits(*5*). We found that, in addition to sleep disruptions, adult flies exhibited circadian arrhythmicity in the setting of pan-neuronal *ISWI* knockdown (**Fig 2A-C**). *ISWI* knockdown led to significantly decreased rest:activity rhythm strength (**Fig 2B**), as well as an increase in the percentage of arrhythmic flies (**Fig 2C; Table S1**). The core molecular clock remained intact (**Fig S3A,B**), suggesting a disruption of clock output mechanisms.

**Fig. 2:**
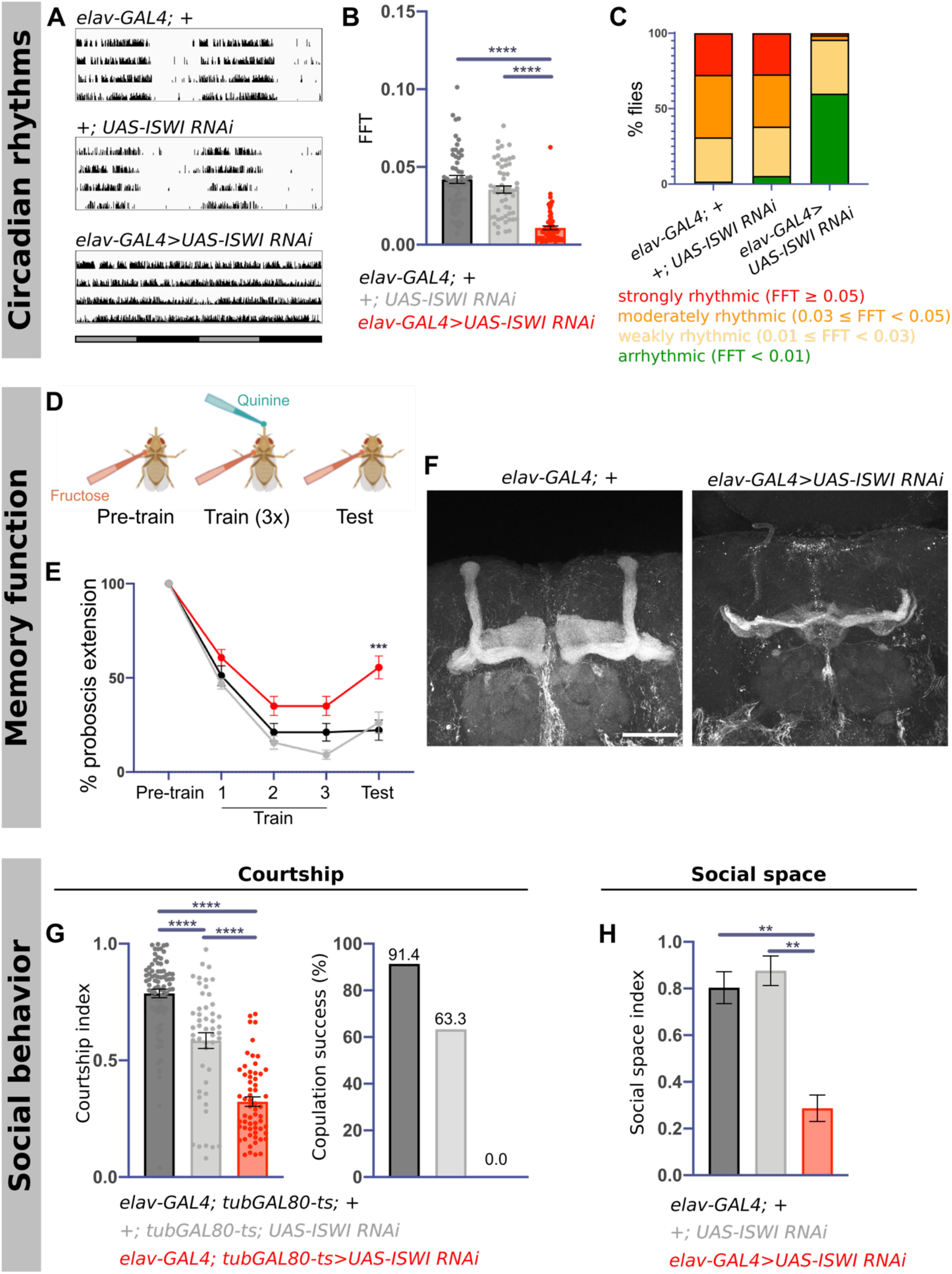
*ISWI* knockdown results in circadian arrhythmicity, memory deficits, and social dysfunction. A) Representative actogram traces from individual flies for each genotype. B) Comparison of rhythm strength in the setting of *ISWI* knockdown (red) and in genetic controls (black and gray), as measured by FFT (n = 58, 55, and 70 from left to right). C) Proportion of strongly rhythmic, moderately rhythmic, weakly rhythmic, and arrhythmic flies in the setting of *ISWI* knockdown and in genetic controls. D) Experimental design of PER. E) Quantification of proboscis extensions in *ISWI* knockdown (blue, n = 39) compared to genetic controls (black, n = 30; gray, n = 47) (Two-way ANOVA with post-hoc multiple comparison test; asterisks denote significance of *ISWI* knockdown condition compared to both genetic controls in post-hoc testing). F) Example FasII staining in controls (left) and *ISWI* knockdown (right). Scale bar, 50um. G) Male courtship indices (left) and copulation success (right) in the setting of *ISWI* knockdown compared to genetic controls (n = 60, 49, 81 from left to right). H) Social space index comparison between *ISWI* knockdown and genetic controls (n ≥ 3 replicates per genotype, 40 flies per replicate per genotype) (see Methods for details on calculation of index).

Since memory disruption is a key characteristic of ID(*32–37*), we next asked whether *ISWI* knockdown leads to memory deficits in adult flies. We assessed aversive taste conditioning using the proboscis extension reflex (PER) assay(*38*) (**Fig 2D**). Flies with pan-neuronal *ISWI* knockdown exhibited intact learning and gustatory responses, as seen by suppressed PER across sequential training sessions; however, these flies erroneously extended their proboscis upon fructose presentation during testing (**Fig 2E**), indicating memory deficits. The mushroom body (MB) is an associative center in the insect brain that is important for normal memory, including conditioning responses seen with PER(*38*). We therefore examined whether ISWI knockdown affects MB structure, and found severe morphologic abnormalities in this brain region: in 100% of *elav-GAL4 > UAS-ISWI RNAi* brains, we observed bilateral ablation of the vertical α/β lobes and thinning of the horizontal γ lobes. (**Fig 2F**). Co-expressing *UAS-ISWI^Res^-FH* with *UAS-ISWI RNAi* was sufficient to rescue MB structure (**Fig S2G**). Together, these results suggest *ISWI* knockdown disrupts MB morphology and memory function. Interestingly, although the MB is known to be involved in adult fly sleep(*39, 40*), we found MB morphologic deficits were dissociable from sleep abnormalities: expression of *ISWI* RNAi using the MB driver *OK107-GAL4* did not alter sleep (**Fig S3C-F)** but disrupted MB morphology (**Fig S3G,H**), suggesting MB dysfunction is not likely to underlie sleep deficits seen in the setting of *ISWI* knockdown.

Social dysfunction is another prevalent symptom in NDDs(*5*). In the male fly, courtship is a social behavior that can be assayed based on a series of stereotyped behaviors. Pan-neuronal *ISWI* knockdown in male flies using the *elav-GAL4* driver was lethal, but restricting knockdown to pre-eclosion using the TARGET system(*41*) (**Fig 3A**) led to viable males. Male flies with *ISWI* knockdown limited to the pre-adult stage exhibited significantly decreased courtship index (time spent courting/total time of assay) and copulation success compared to genetic controls (**Fig 2G**), suggesting compromised social function. To provide further evidence that *ISWI* knockdown affects social rather than only reproductive behaviors, we utilized a social space behavioral assay in which distance between individual flies is measured in a two-dimensional space(*42*). Indeed, female *elav-GAL4>UAS-ISWI RNAi* flies exhibited increased social space in relation to genetic controls (**Fig 2H; Fig S3I,J**). This result supports the conclusion that social behaviors are disrupted with *ISWI* knockdown, independent of reproductive function. Thus, a *Drosophila* homolog of an NDD-associated human gene is required for normal sleep, circadian rhythmicity, memory, and social behaviors.

**Fig. 3:**
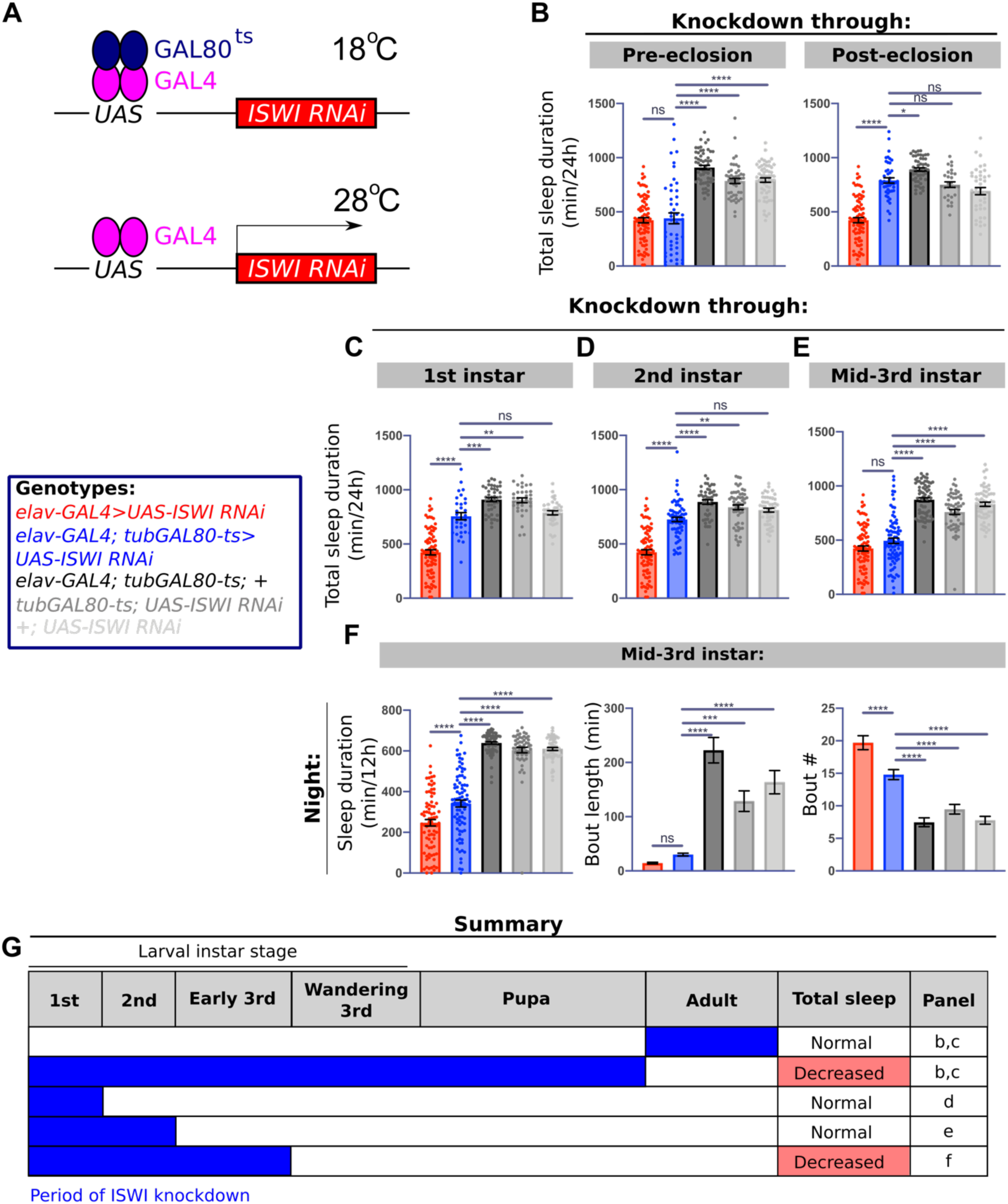
*ISWI* knockdown through mid-3^rd^ instar leads to sleep disruptions. A) Schematic of TARGET system. B) Quantification of total sleep duration in the setting of pre- (left) or post-eclosion (right) *ISWI* knockdown (blue) compared to genetic controls (black and grays) exposed to the same temperature shifts across development or constitutive knockdown (red) (always at 25°C) (for pre-eclosion: n = 82, 46, 57, 47, 58 from left to right; for post-eclosion: n = 82, 46, 52, 28, 40). Quantification of total sleep duration in the setting of *ISWI* knockdown through C) 1^st^ instar (n = 82, 31, 43, 33, 36, from left to right) and D) 2^nd^ instar (n = 82, 69, 46, 57, 50, from left to right). E) Total sleep duration and F) night sleep duration, sleep bout length, and sleep bout number (left to right) in the setting of *ISWI* knockdown through mid-3^rd^ instar (n = 82, 84, 68, 55, 58, from left to right). G) Summary of *ISWI* knockdown periods and resulting effects on total sleep duration.

### ISWI is required during the 3^rd^ instar larval stage for normal adult sleep

*ISWI* and its homologs are involved in neural development and differentiation across species(*13, 16–18*). We asked whether *ISWI* is required during pre-adult developmental stages or in an ongoing manner in the adult fly to regulate adult behaviors. We leveraged the TARGET system(*43*) to restrict *ISWI* knockdown to pre- or post-eclosion (**Fig 3A**). Pan-neuronal knockdown only during pre-eclosion significantly decreased total sleep and resulted in sleep fragmentation, recapitulating the phenotype seen with constitutive *ISWI* knockdown (**Fig 3B; Fig S3A-D**). In contrast, knockdown restricted to the adult had no effect on total sleep or sleep fragmentation (**Fig 3B; Fig S3A-D**). More refined temporal mapping revealed pan-neuronal *ISWI* loss from embryonic stages through the mid-3^rd^ instar period leads to decreased total sleep duration and sleep fragmentation similar to constitutive ISWI knockdown (**Fig 3E-G; Fig S3A-D**), whereas knockdown only through earlier stages does not (**Fig 3C,D,G; Fig S3A-D)**.

Given the pre-eclosion role of *ISWI* in determining adult sleep, we wondered whether sleep deficits arise prior to adulthood. We previously characterized sleep behaviors during the 2^nd^ instar larval stage(*44*), and found here that *ISWI* knockdown has no effect on sleep during this larval period (**Fig S4E**). This result is consistent with our finding that *ISWI* knockdown through 2^nd^ instar does not affect adult sleep (**Fig 3D**), and is likely acting during mid-3^rd^ instar to affect adult sleep behaviors, presumably by coordinating development of adult sleep-regulatory circuits.

Less severe effects on total sleep duration and sleep fragmentation were also observed with knockdown through 1^st^ and 2^nd^ instar stages (**Fig 3C,D; Fig S3A-D**). These phenotypes did not fully recapitulate constitutive *ISWI* loss, leading us to ask whether such sleep changes were secondary to circadian disturbances. Temporal mapping of rest:activity rhythm defects revealed pre-eclosion *ISWI* knockdown also led to adult arrhythmicity (**Fig 4A-C**). However, in contrast to sleep, knockdown through only the 1^st^ instar stage led to adult rest:activity arrhythmicity comparable to constitutive ISWI knockdown (**Fig 4A; Fig S5; Table S2**). This result indicates that *ISWI* knockdown through earlier larval stages primarily results in adult rhythmic deficits, associated redistribution of sleep across the 24-hour day, and sleep fragmentation. Taken together, our findings suggest that primary sleep disruptions and behavioral arrhythmicity are temporally separable to distinct developmental windows of *ISWI* knockdown.

**Fig. 4:**
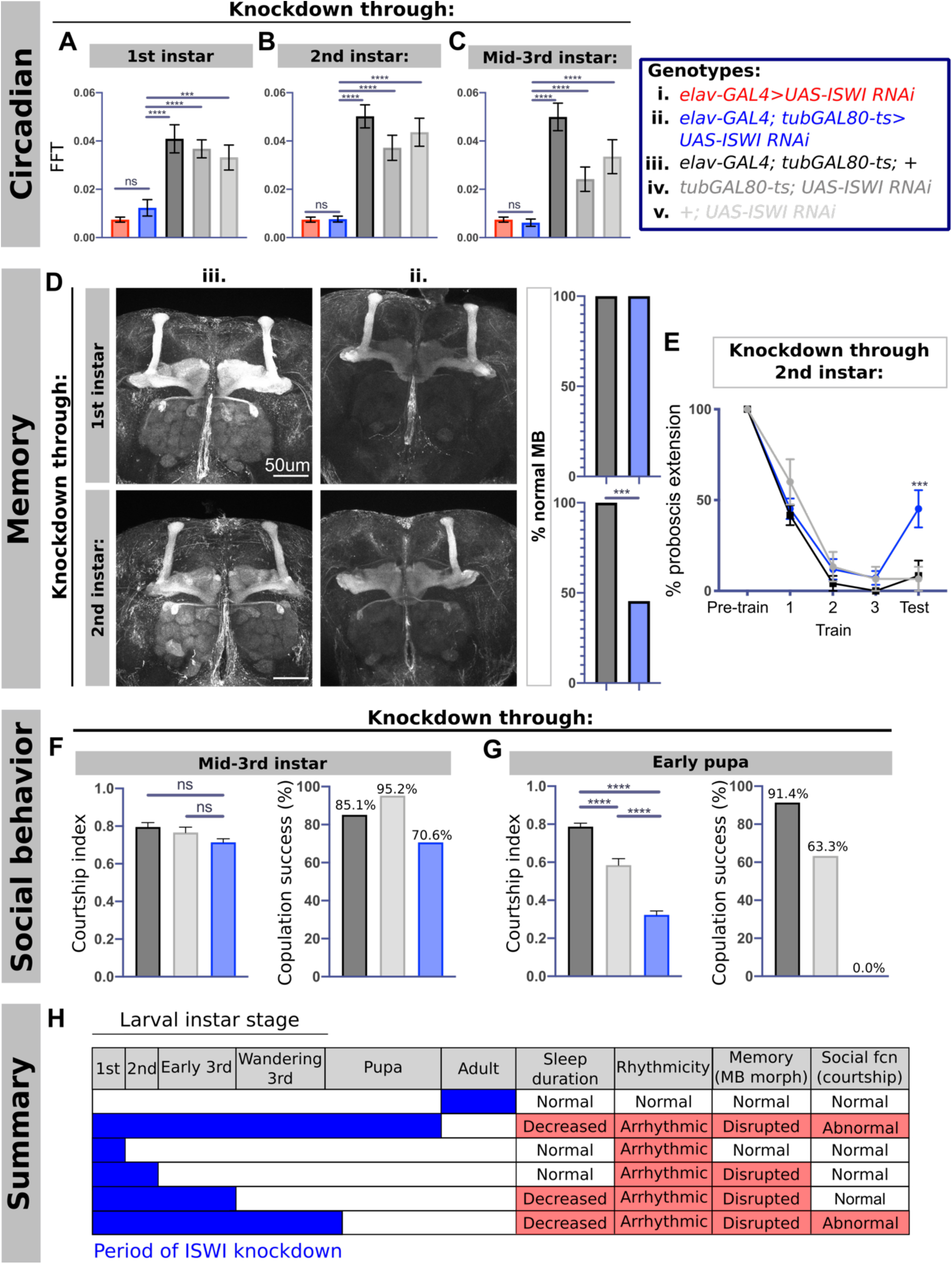
*ISWI* knockdown during separable developmental windows leads to distinct behavioral phenotypes. A) Quantification of rhythmicity as measured by maximum FFT amplitude of temporally restricted *ISWI* knockdown (blue) through B) 1^st^, C) 2^nd^, and D) mid-3^rd^ instars (left to right) compared to constitutive knockdown (red) and genetic controls (black and grays) (for all *elav* > UAS-ISWI RNAi (red), n = 23; from left to right for all other conditions: knockdown through 1^st^ instar, n = 25, 28, 27, 27; knockdown through 2^nd^ instar, n = 30, 32, 30, 30; knockdown through mid-3^rd^ instar, n = 29, 31, 30, 18). D) Representative images of FasII immunostaining with *ISWI* knockdown through 1^st^ (top) and 2^nd^ instars (bottom), with quantification of percentage of brains with normal MB morphology (for knockdown through 1^st^ instar: n = 12, black; n = 15, blue; for knockdown through 2^nd^ instar: n = 21, black; n = 22, blue) (Fisher’s Exact test). E) Quantification of PER assay with temporally-restricted *ISWI* knockdown through 2^nd^ instar (blue; n = 14) compared to genetic controls (black, n = 10 and gray, n = 12) (Two-way ANOVA with post-hoc multiple comparison test; asterisks denote significance of *ISWI* knockdown condition compared to both genetic controls in post-hoc testing). (F, G) Courtship index (left) and copulation success (right) for *ISWI* knockdown through F) mid-3^rd^ (n = 39, 62, 42 from left to right) and G) early pupation (n = 60, 81, 49 from left to right). H) Summary of windows of *ISWI* knockdown that give rise to different adult phenotypes.

### ISWI acts during separable pre-adult stages for adult fly memory and social functions

We next investigated the temporal window of ISWI action for adult memory and courtship behaviors. In contrast to sleep, knockdown through only the mid-2^nd^ instar stage led to MB morphologic abnormalities and deficits in aversive taste conditioning (**Fig 4D,E**). These results suggest the ISWI-dependent sleep phenotype does not arise from MB disruptions, as sleep and memory deficits are temporally dissociable. This conclusion is further supported by our finding that *ISWI* knockdown in the MB (using *OK107-GAL4*) is associated with MB but not sleep deficits (**Fig S3C-H**), underscoring *ISWI* functions in distinct circuits and developmental times for sleep and memory. Finally, male flies exhibited disrupted courtship behavior with *ISWI* knockdown through early pupation, but not through mid-3^rd^ instar (**Fig 4F,G**), dissociating sleep and courtship behaviors. These results demonstrate *ISWI* acts in different developmental windows to coordinate distinct adult behaviors (**Fig 4H**).

### ISWI function in type I neuroblasts is necessary for normal adult sleep and MB morphology

How does *ISWI* affect development of adult fly sleep circuits? Since *elav* is expressed pan-neuronally(*45*), we reasoned *ISWI* loss in specific neurons might result in adult fly sleep deficits. We performed a large neuronal GAL4 screen (>400 lines), but found none of the tested lines recapitulated sleep loss seen with *elav-GAL4 > UAS-ISWI RNAi* (data not shown). These negative results led us to wonder whether the sleep phenotype seen with *elav-*driven *ISWI* knockdown is not related to *ISWI* function in neurons. *Elav* is also expressed in larval glial cells and neuroblasts(*46*), but restricting *ISWI* knockdown to glia using *repo-GAL4* had no effect on sleep (**Fig S6A,B**), arguing against a glial role. *ISWI* is necessary for maintaining chromatin structure in larval neuroblasts(*13, 14*) and normal progenitor cell proliferation across species(*15, 18*). To test whether *ISWI* RNAi in all neuroblasts leads to sleep deficits, we knocked down *ISWI* using *worniu-GAL4* and observed a reduction in sleep duration (**Fig 5A,B**). In addition, *worniu-GAL4 > UAS-ISWI RNAi* flies exhibited decreased night-time sleep and bout length, comparable to *elav-GAL4* > *UAS-ISWI RNAi* flies (**Fig 5C,D**). Knocking down *ISWI* in all neuroblasts also led to disruptions in MB morphology (100% of MB were abnormal in *worniu-GAL4 > UAS-ISWI RNAi* flies) similar to that seen with *elav-GAL4* driven knockdown (**Fig 5E**). Consistent with the hypothesis that *ISWI* functions in neuroblasts, *OK107-GAL4* expresses in MB neuroblasts(*47*) and also disrupts MB morphology with *ISWI* knockdown (**Fig S3G,H**).

**Fig. 5:**
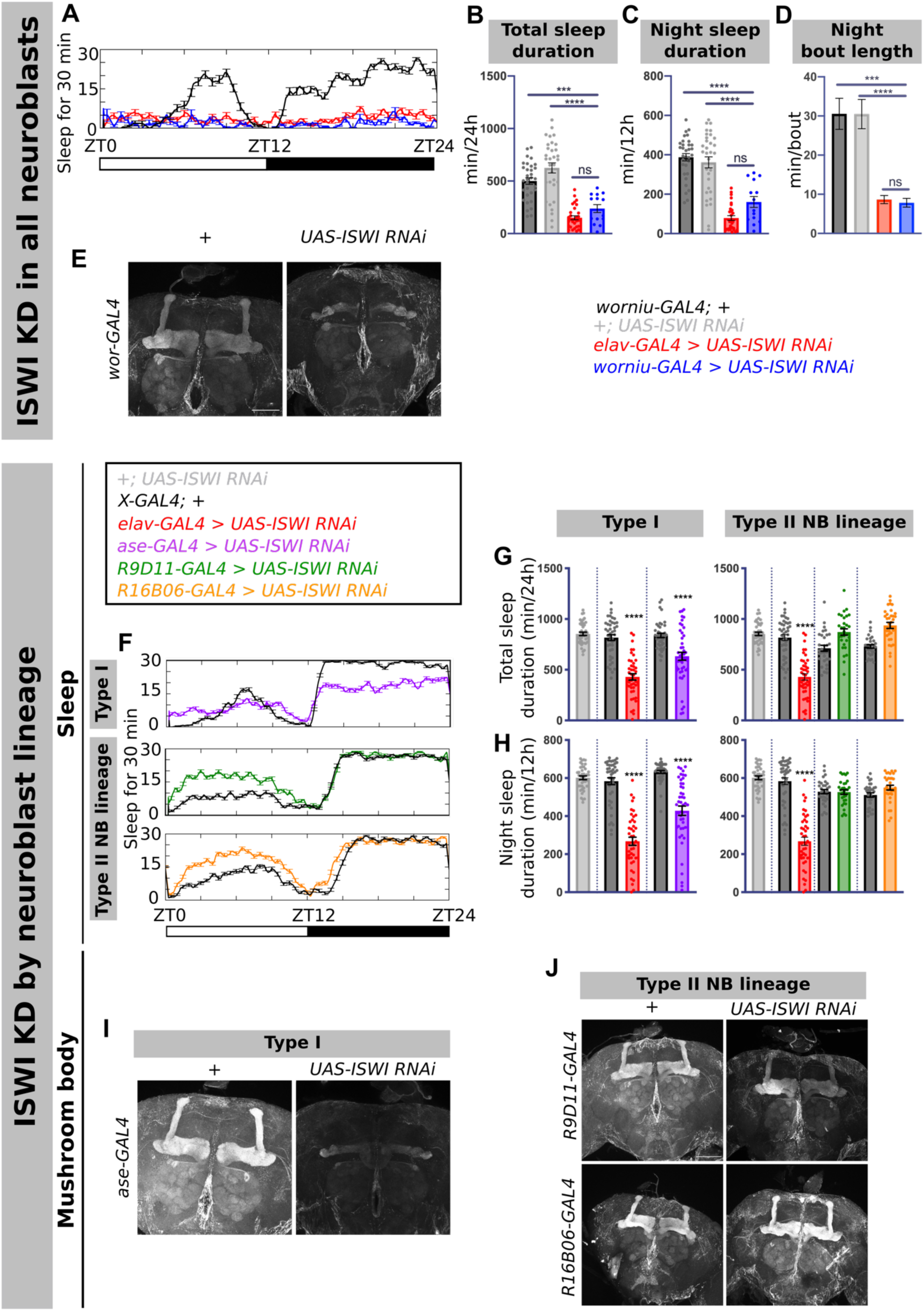
*ISWI* knockdown in type I neuroblasts leads to sleep disruption and MB morphologic deficits. A) Representative sleep traces from multibeam sleep monitoring with *ISWI* knockdown in all neuroblasts using *worniu-GAL4* driver (blue) compared to *elav-*driven knockdown (red) and genetic controls (black and gray). Quantification of B) total sleep duration, C) night sleep duration, and D) night sleep bout length in *worniu-GAL4 > UAS-ISWI RNAi* flies as measured by multibeam monitoring compared to *elav-GAL4 > UAS-ISWI RNAi* and genetic controls (n = 32, 32, 31, 14 from left to right). E) Representative images of FasII immunostaining of brains of *worniu-GAL4 > UAS-ISWI RNAi* flies. F) Representative sleep traces for *ISWI* knockdown in different neuroblast lineages. Quantification of G) total sleep duration and H) nighttime sleep with *ISWI* knockdown in different neuroblast lineages (from left to right: type I neuroblasts, n = 42, 43, 43, 42, 47; type II neuroblast lineages, n =42, 43, 43, 31, 30, 31, 32. Asterisks denote significance compared to both the *GAL4* only control and *UAS-ISWI RNAi* only control). Representative images of FasII immunostaining of brains with *ISWI* knockdown in I) type I neuroblasts and J) type II neuroblast lineages.

We next asked whether *ISWI* knockdown in specific neuroblast lineages are responsible for adult sleep and MB deficits. In the developing *Drosophila* nervous system, type I and II neuroblasts undergo asymmetric cell divisions; type II divide into intermediate progenitor cells (INPs), which are capable of several rounds of cell division before differentiating into neurons(*48, 49*). *ISWI* knockdown in type I neuroblasts using *asense-GAL4* significantly decreased sleep duration (**Fig 5F-H**) and increased sleep fragmentation (**Fig S6C,D**). *Asense* is also expressed in type II lineage INPs(*49*), so the sleep phenotype with *asense-GAL4* could result from *ISWI* knockdown in INPs rather than type I neuroblasts. However, knockdown in INPs using *R9D11-GAL4* or *R16B06-GAL4*(*50*) did not disrupt sleep (**Fig 5F-H; Fig S6C,D**), suggesting ISWI acts in type I neuroblasts for adult fly sleep behaviors. Similarly, *ISWI* knockdown in type I neuroblasts, but not in INPs, resulted in MB morphologic deficits (**Fig 5I,J**). Thus, *ISWI* function in type I neuroblasts during development is required for normal adult sleep and MB structure.

### ISWI knockdown disrupts morphology and function of sleep-regulatory dorsal fan-shaped body neurons

To understand how *ISWI* knockdown affects sleep regulatory circuits, we focused on the adult sleep-promoting dorsal fan-shaped body (dFB) neurons that are defined by the *R23E10* enhancer. dFB neurons are involved in the homeostatic sleep response: sleep deprivation increases activity of dFB neurons, and activation of these neurons induces sleep(*51–53*). Since *ISWI* knockdown results in sleep rebound deficits following mechanical deprivation (**Fig 1J; Fig S1D-F**), we hypothesized the sleep-promoting function of dFB neurons might be impaired with *ISWI* knockdown. In the setting of pan-neuronal *ISWI* knockdown, *R23E10* adult neurons exhibited abnormal neurite morphology, with aberrant projections to brain regions outside of the dFB; we also observed abnormal cell body location of *R23E10* adult neurons in the context of *ISWI* knockdown compared to genetic controls (**Fig 6A**). Quantification of *R23E10* axonal innervation of the dFB showed decreased innervation volume in the setting of *ISWI* knockdown (**Fig 6B**). In addition, there was a significant increase in the total number of *R23E10* neurons as measured by GFP+ soma (**Fig 6C**), suggesting *ISWI* knockdown disrupts dFB neuron cell fate. Of note, *R23E10-GAL4>UAS-ISWI RNAi* adult flies showed no sleep changes (**Fig S7A,B**). Since ISWI is required during the mid-3^rd^ instar stage for normal adult sleep, this result was anticipated because the R23E10 driver does not label primordial dFB neurons in the larval nervous system(*54*).

**Fig. 6:**
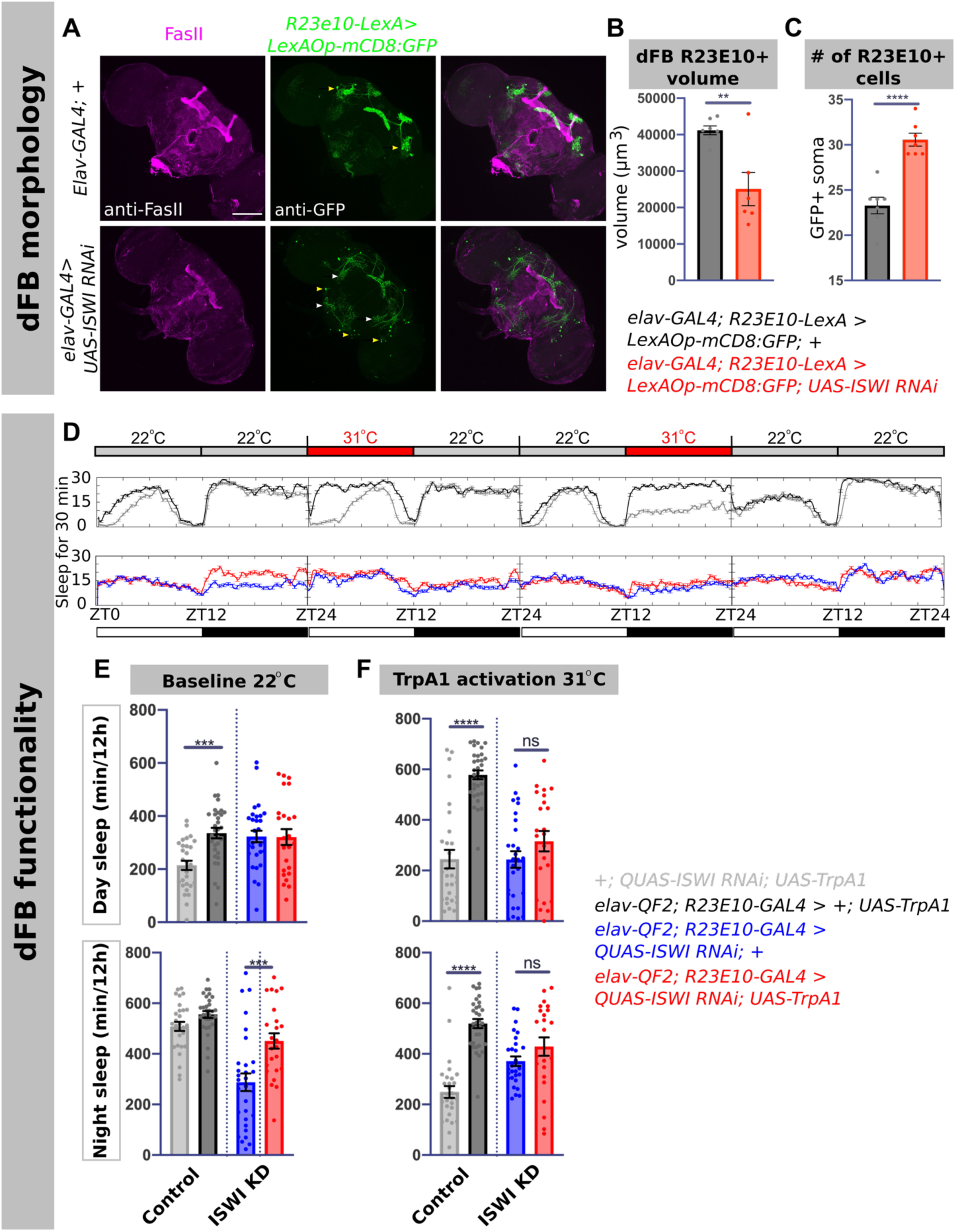
*ISWI* knockdown disrupts the morphology and function of the sleep-promoting dFB. A) Representative images of *R23E10* neuron morphology as visualized by GFP staining (middle panels) in genetic controls (top) and in the setting of *ISWI* knockdown (bottom), with FasII counterstaining (left panels). White arrowheads point to abnormal *R23E10* neuron projections, yellow arrowheads indicate normal and aberrant cell body locations in genetic controls and in the setting of *ISWI* knockdown, respectively. Scale bar, 100um. Quantification of B) dFB volume and C) number of *R23E10* soma in genetic controls (black, n = 10) and in the setting of pan-neuronal *ISWI* knockdown (red, n = 15) as measured by GFP immunostaining. D) Thermogenetic activation of *R23E10* neurons, with experimental design showing temperature shifts (top) and representative sleep traces (bottom panels). Quantification of day (top) and night (bottom) sleep at E) 22°C baseline and F) sleep in the setting of TrpA1 activation at 31°C across all experimental and control groups (n = 32, 29, 24, 30 from left to right).

We next asked whether temporally restricting *ISWI* knockdown to the early developmental window associated with adult sleep deficits would affect *R23E10* neuron morphology. In the setting of *ISWI* knockdown through mid-3^rd^ instar, we found decreased dFB innervation volume and increased cell body number **(Fig S7C**). In contrast, while we observed MB morphologic deficits with *ISWI* knockdown restricted through 2^nd^ instar (**Fig 4D**), dFB morphology remained unchanged compared to genetic controls **(Fig S7C**). These results parallel our earlier findings that *ISWI* knockdown through 2^nd^ instar does not affect adult sleep, while knockdown through mid-3^rd^ instar results in adult sleep deficits. Thus, *ISWI* knockdown appears to disrupt development of a key group of sleep-promoting neurons, leading to adult sleep deficits.

Since we observed that both dFB morphologic changes and behavioral sleep abnormalities mapped to the same developmental window, we next asked whether *ISWI* knockdown resulted in functional dFB abnormalities that could underlie the sleep phenotype. In the setting of *elav-*driven *ISWI* knockdown, we thermogenetically activated *R23E10* cells by expressing TrpA1, a heat-sensitive cation channel(*55*). We collected 24 hours of baseline sleep (**Fig 6D,E**) and then activated *R23E10* neurons during the day or night (**Fig 6D,F**). While genetic controls exhibited a robust increase in sleep at the activation temperature during either day or night, flies with *ISWI* loss failed to show this change (**Fig 6F**). To address differences in baseline sleep between genetic conditions (**Fig 6E**), we also measured sleep differences between activation and baseline periods in individual flies (**Fig S7D**). Using this measurement, we found again that *ISWI* knockdown attenuated the functional effects of *R23E10* activation (**Fig S7E**). Together, these results suggest *ISWI* knockdown disrupts the development of dFB neurons and impairs their sleep-promoting function.

### SMARCA1 and SMARCA5 differentially rescue adult sleep and MB defects

We next tested whether wild-type human *SMARCA1* (*SMARCA1^WT^*) and *SMARCA5* (*SMARCA5^WT^*) could rescue adult sleep and MB defects in the setting of ISWI knockdown. Driving *UAS-SMARCA1^WT^* expression with *elav-*GAL4 did not rescue ISWI knockdown-induced sleep deficits (**Fig 7A,B**), but did restore MB morphology as measured by vertical and horizontal lobe volumes (**Fig 7D,E**). Conversely, *SMARCA5* expression restored sleep (**Fig 7A,B; Fig S8**) but not all aspects of MB morphology (**Fig 7D,E**). Notably, driving *SMARCA5* expression did not decrease activity index in the setting of *ISWI* knockdown (**Fig 7C**), demonstrating that sleep rescue with *SMARCA5* co-expression was not confounded by impaired locomotor behavior. These results suggest the human homologs of ISWI act separately for the development of circuits involved in different behaviors.

**Fig. 7:**
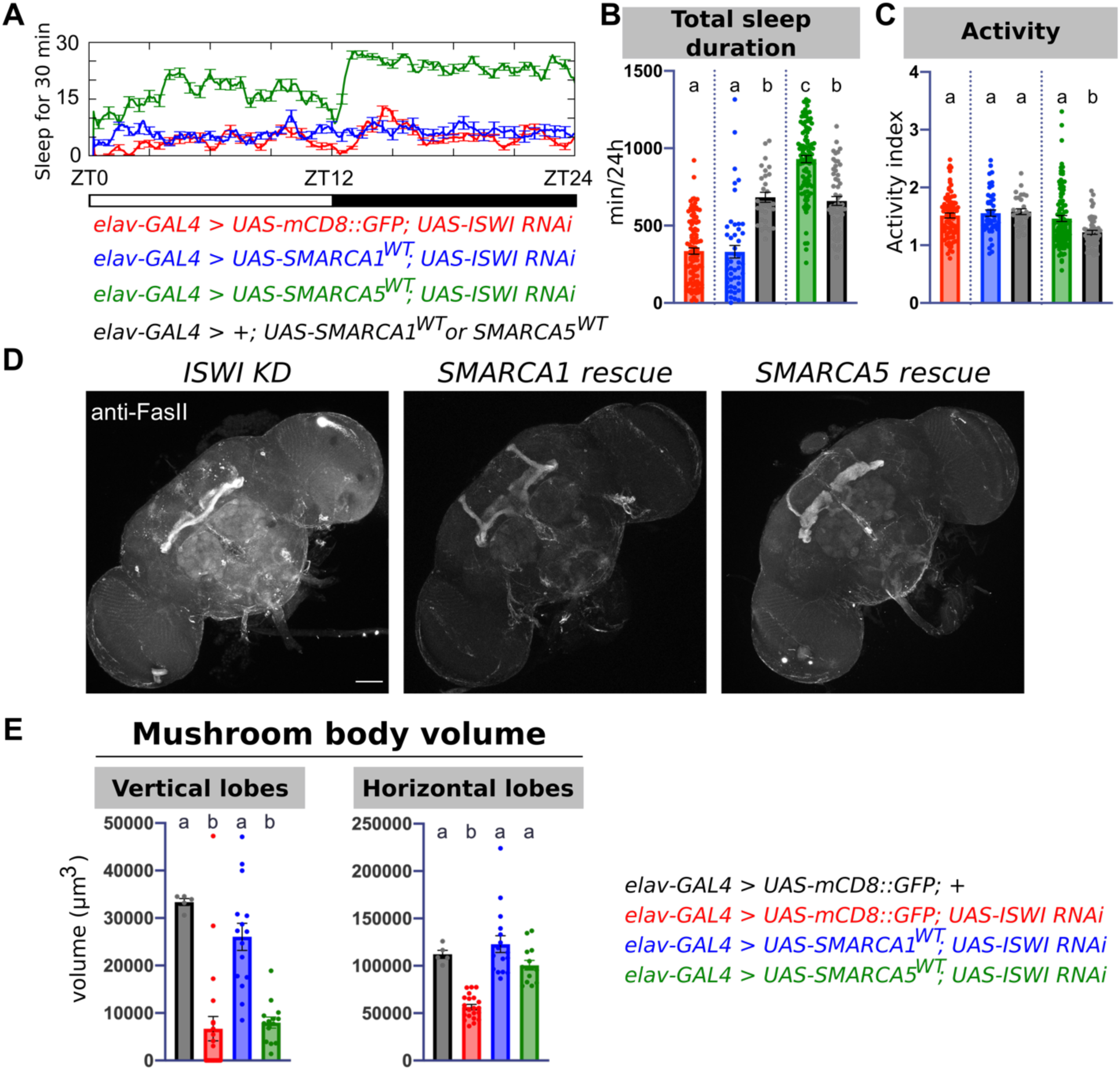
The human *ISWI* homologs *SMARCA1* and *SMARCA5* differentially rescue sleep and MB morphology in the setting of *ISWI* knockdown. A) Representative sleep traces with *ISWI* knockdown (red) compared to *SMARCA1^WT^* (blue) and *SMARCA5^WT^* (green) expression in the setting of *ISWI* knockdown, or overexpression alone (black) with B) quantification of total sleep and C) activity index across experimental and control groups (n = 104, 48, 29, 102, 55 from left to right). D) Representative images of FasII immunostaining of adult brains with pan-neuronal *ISWI* knockdown (left), *SMARCA1^WT^* rescue (middle), and *SMARCA5^WT^* rescue (right), and E) quantification of MB morphology across groups. Volumes are presented as the sum of horizontal (left) or vertical (right) lobe volumes by each brain (n = 5, 21, 16, 14 from left to right).

To further investigate mechanisms by which *ISWI* and *SMARCA5* regulate sleep, we performed RNA-Seq analysis on mid-3^rd^ instar larval central nervous systems (CNS) in the setting of pan-neuronal ISWI knockdown (**Fig S9A**). We chose mid-3^rd^ instars because temporal mapping revealed *ISWI* knockdown during this developmental stage disrupts adult *Drosophila* sleep (**Fig 3E,F**). Differential gene expression analysis showed ISWI knockdown resulted in 687 differentially expressed genes (DEGs; 381 downregulated and 306 upregulated genes) (**Fig S9B**). These DEGs were enriched for genes involved in several neuronal and cellular functions, including axon guidance, mesoderm development, and cell adhesion. Using 679 human homologs of these DEGs, we performed connectivity analysis in the context of a brain-specific gene interaction network(*56, 57*). Compared to all genes in the network, *SMARCA5* exhibited significantly enhanced connectivity to human homologs of *Drosophila* DEGs (**Fig S9C, D**), suggesting high concordance between *ISWI* and *SMARCA5* in their gene interaction networks. *SMARCA1* also exhibited significantly increased connectivity to human DEG homologs (**Fig S9D**). *SMARCA5* additionally exhibited increased connectivity to 224 human genes involved in sleep and circadian rhythm functions (**Fig S9D**) compared to all genes in the network. The “connector genes” located in the shortest paths between *SMARCA5* or *SMARCA1* to DEGs in the network were enriched for basic cellular processes such as gene expression, chromosome organization, response to stress, and DNA repair (**Fig. S9E**). Moreover, connector genes between *SMARCA5* and human sleep-related genes were involved in neurogenesis, nervous system development, and Wnt signaling pathways. These results underscore the relevance of *ISWI* in sleep function and neurodevelopment, and implicate a developmental role for *SMARCA5* in human sleep gene networks.

## Discussion

Despite clinical heterogeneity among and even within NDDs, sleep disturbances are highly prevalent across these diverse disorders(*3, 4*). Clinical evidence points towards a link between sleep dysfunction and other behavioral symptoms in NDDs(*5, 6, 58*). Whether sleep disturbances are a byproduct of other NDD-related deficits or directly result from developmental disruptions remains a point of debate(*3, 7–9*). From a sleep-focused screen of *Drosophila* orthologs of human NDD-associated genes, we found that the chromatin remodeler *ISWI* is important for adult fly sleep. Knockdown in distinct developmental windows and circuits resulted in dissociable adult deficits in sleep, circadian, memory, or social behaviors. Notably, along with other behavioral deficits, our findings demonstrate that sleep disruptions represent a primary phenotype arising directly from *ISWI* knockdown during pre-adult development.

Mutations in chromatin remodelers are strongly associated with NDDs. For example, de novo mutations in the chromatin remodeler *CHD8* have the strongest overall association with autism spectrum disorders (ASDs)(*22*). Indeed, in addition to its role in growth-regulatory pathways(*59, 60*), *CHD8* has been shown to interact with and control the expression of other autism risk genes(*61, 62*). These lines of evidence suggest chromatin remodelers are ideal candidates to identify the mechanisms by which behavioral pleiotropy arises in NDDs. Our results trace how disruptions in a single NDD risk gene affects development of neural circuits controlling distinct behaviors, and identify a genetic etiology underlying NDD-associated sleep disruption.

*ISWI* and its homologs have been implicated in neural stem cell fate decision, and *ISWI* is necessary for proper chromatin regulation in larval neuroblasts(*13–15*). Mouse models with mutations in *Smarca1* and *Smarca5* exhibit abnormal neural progenitor cell proliferation(*16–18*). In our present study, we have begun to parse out the stem cell lineage and timing of events that lead to specific disruptions in adult behaviors. *ISWI* knockdown in type I neuroblasts disrupted adult sleep, and resulted in disrupted morphology and function of sleep-promoting *R23E10* neurons, suggestive of abnormalities in neural stem cell proliferation and differentiation. These results raise the possibility that *ISWI* knockdown changes the fate of adult sleep-regulatory neurons, perhaps through dysregulation of temporally expressed transcription factors or cellular signaling important for neuroblast differentiation. We also found *ISWI* function in type I neuroblasts to be important for normal MB morphology. Notably, knockdown in MB neuroblasts, which are type I neuroblasts, was sufficient to disrupt MB morphology but not sleep, distinguishing the neural substrates underlying *ISWI-*related sleep and memory deficits. Together with our temporal mapping results, we propose that *ISWI* affects the fate of neural stem cells contributing to adult circuits responsible for separate adult behaviors during the course of larval development. These results demonstrate the importance of ISWI chromatin remodelers for development of normal adult behaviors, building on existing evidence that *ISWI* plays a critical role in neural stem cell differentiation. Determining which populations of stem cells are affected at a given stage of pre-adult development, and tracing how *ISWI* knockdown affects formation of specific circuits, is the next step towards understanding how *ISWI* loss disrupts adult behaviors.

It remains unknown whether *ISWI* loss specifically in *23E10* cells is causative for the observed sleep deficits. One limitation of the GAL4/UAS system is the shifting expression patterns of GAL4 drivers across development(*54, 63*): the *23E10* driver labels sleep-promoting dFB neurons in the adult fly, but is expressed in different cells at mid-3^rd^ instar. Congruent with this, *ISWI* knockdown with *23E10* has no effect on adult fly sleep. Work is needed to identify and genetically access the relevant primordial sleep cells in the larval nervous system.

We found that the human homologs of *ISWI, SMARCA1* and *SMARCA5*, are able to rescue sleep deficits and MB disruptions, respectively. Why do *SMARCA1* and *SMARCA5* differentially rescue sleep and MB abnormalities? One possible explanation lies in mouse studies that have noted differences in temporal and spatial distributions of *Smarca1* and *Smarca5* in the developing mouse brain. Differences in protein sequences between SMARCA1 and SMARCA5 may also facilitate differential expression and function. Our results begin to parse the differential functions of *SMARCA1* and *SMARCA5*, an outstanding question in the field. This is an area of major interest, as understanding the differences between paralogs in the setting of disease can inform our knowledge about compensatory effects that paralogs may exert to alter phenotypes.

Although mutations in the human *ISWI* homolog *SMARCA1* have been implicated in diverse NDDs(*19–21*), to date, there has been no clinical characterization of sleep phenotypes arising from patient mutations in *SMARCA1* and *SMARCA5*. A human brain-specific gene network analysis shows that *ISWI* and its human homologs interact with a conserved network of genes. Compellingly, *SMARCA5,* which rescued sleep deficits in the setting of *ISWI* knockdown, exhibited increased connectivity to human sleep and circadian genes through connector genes broadly involved in development. These results implicate a role for *SMARCA5* in the development of normal human sleep regulation. It will be of great interest to assess sleep in patients with *SMARCA5* mutations given our findings in flies. Moreover, because *ISWI* knockdown leads to sleep abnormalities in the adult fly, longitudinal patient sleep phenotyping may reveal sleep differences across the lifespan. In sum, our results provide new insight into the etiology of sleep disruptions in NDDs, and suggest a mechanism whereby temporally and spatially constrained gene function underlies behavioral pleiotropy. Importantly, this work supports the idea that sleep is a developmentally-programmed behavior; sleep abnormalities in NDDs are not simply a byproduct of broad cognitive/behavioral deficits, but rather emerge from specific developmental anomalies.

## Materials and Methods

### Fly stocks

Flies were raised and maintained on standard molasses food (8.0% molasses, 0.55% agar, 0.2% Tegosept, 0.5% propionic acid) at 25°C on a 12hr:12hr light:dark cycle unless otherwise specified. Unless otherwise specified, female flies were used in all experiments.

### Fly strains

The *hs-hid; elav-GAL4; UAS-Dcr2* strain was a gift of Dr. Dragana Rogulja (Harvard University). *Elav^C155^-GAL4 (elav-GAL4), OK107-GAL4, repo-GAL4, 23E10-GAL4,* were gifts of Dr. Amita Seghal (University of Pennsylvania). *Worniu-GAL4, asense-GAL4, and worniu-GAL4, ase-GAL80* were gifts of Dr. Mubarak Syed (University of New Mexico). *UAS-dTrpA1* was a gift from Dr. Leslie Griffith (Brandeis University). The following strains were purchased from the Bloomington Drosophila Resource Center: *UAS-ISWI RNAi^HMS00628^* (*UAS-ISWI RNAi^TRiP^*) was used for all experiments unless otherwise specified and from the Harvard Transgenic RNAi Project (TRiP) (BSC #32845); *UAS-mCD8::GFP* (BSC #5137); *tub-GAL80^ts^* (BSC #7019); *23E10-GAL4* (*BSC #49032*)*; LexAOp-mCD8::GFP* (BSC #32203); *R9D11-GAL4 (BSC #40731); R16B06-GAL4 (BSC # 40731); elav^C155^-QF2* (*elav-QF2;* BSC #66466). All RNAi strains used in the primary screen were purchased from Bloomington Drosophila Resource Center (see Table S1 for a full list of lines). *UAS-ISWI RNAi^GD1467^* (*UAS-ISWI RNAi^VDRC^*) was purchased from the Vienna Drosophila Resource Center (VDRC #24505, construct ID GD1467). The following fly strains were generated as described below:

<u>UAS-ISWI^Res^-FH construct:</u> A vector containing the ISWI gene sequence with a C-terminal FLAG-HA tag under UAS control (UFO10052) was obtained from the Drosophila Genomics Resource Center (NIH Grant 2P40OD010949). The gene location targeted by the HMS00628 ISWI RNAi hairpin (5’ ACCCAAGAAGATCAAAGACAA 3’) was identified. The Q5 Site-Directed Mutagenesis Kit (New England BioLabs, cat#E0554S) and corresponding primer design tool were used to create RNAi-resistant UAS-ISWI. Primers were as follows:

- Forward: ttaaggataaGGACAAGGAAAAGGATGTG
- Reverse: tttttttaggcCTACCCTTAGGCTTCGTG.

DNA injection was prepared with the Midiprep Kit (Qiagen). Injections were performed by Rainbow Transgenic Flies, Inc for production of transgenic flies at the attP40 landing site.

<u>QUAS-ISWI RNAi construct:</u> QUAS-WALIUM20 vector was obtained from J. Zirin at the Fly Transgenic RNAi Project(*64*). The HMS00628 ISWI RNAi hairpin, originally used to generate the UAS-ISWI RNAi construct (BSC #32845), was cloned into the QUAS-WALIUM20 vector using the pWALIUM20 cloning protocol (available at www.flyrnai.org). Briefly, the following oligonucleotides were synthesized and annealed (21 NT hairpin sequence shown in capital letters):

- 5’ctagcagtACCCAAGAAGATCAAAGACAAtagttatattcaagcataTTGTCTTTGATCTTCTTGGGTgcg 3’
- 5’aattcgcACCCAAGAAGATCAAAGACAAtatgcttgaatataactaTTGTCTTTGATCTTCTTGGGTactg 3’.

The QUAS-WALIUM20 vector was linearized by NheI and EcoRI, and the DNA fragment containing the hairpin was ligated into the vector. DNA injection was prepared with the Midiprep Kit (Qiagen). Injections were performed by Rainbow Transgenic Flies, Inc for production of transgenic flies at the attP40 and VK00033 landing sites.

<u>UAS-SMARCA1 construct:</u> Flies carrying UAS-SMARCA1^WT^ were generated using human *SMARCA1* cloned into the pFastBac Dual vector (Addgene plasmid #102243). Gateway cloning (Invitrogen) was used to generate the pDonr221-20xUAS (a gift from Dr. Paula Haynes) and pDonrP2rP3-SMARCA1. Primers for pDonrP2rP3-SMARCA1 were designed by fusing Gateway attB2r and attB3 sequences upstream and downstream, respectively, of the SMARCA1 sequence. Primer sequences were as follows (capitalized letters indicate Gateway sequences):

- Forward, attB2r-SMARCA1: GGGGACAGCTTTCTTGTACAAAGTGGatggaacaagacactgctgcc
- Reverse, attB3-SMARCA1: GGGGACAACTTTGTATAATAAAGTTGttacgacttcaccttcttcacatc.

A modified pBPGUw, pBPGUw-R1R3-p10(*65*) (a gift from Dr. Paula Haynes) was used for gateway recombination. Injections were performed by Rainbow Transgenic flies, Inc for production of transgenic flies at the attP40 landing site.

<u>UAS-SMARCA5 construct</u>: The coding sequence of SMARCA5^WT^ was synthesized and cloned into the pACU2 vector (GenScript). Injections were performed by Rainbow Transgenic Flies, Inc for production of transgenic flies at the attP2 landing site.

### Sleep assays

Adult female flies were collected 2-3 days post-eclosion and aged in group housing on standard food at 25°C on a 12hr:12hr LD cycle (unless otherwise noted). Flies aged 5-7 days were anesthetized on CO_2_ pads (Genesee Scientific Cat #59-114) and loaded into individual glass tubes (with 5% sucrose and 2% agar) for monitoring locomotor activity in the Drosophila Activity Monitoring (DAM) system (Trikinetics, Waltham MA) or MultiBeam Activity Monitors (Trikinetics, Waltham MA) as denoted in figure legends. All sleep experiments were loaded between ZT5-ZT10. Data collection began at ZT0 at least 24 hours following CO_2_ anesthesia. Activity was measured in 1 minute bins and sleep was defined as 5 minutes of consolidated inactivity(*25, 26*). Data processing was performed using PySolo(*66*). Mechanical sleep deprivation was performed using a Trikinetics vortexer mounting plate. Monitors were shaken for 2 seconds randomly within every 20 second window for 12 hours during the night. Rebound sleep was calculated as the difference between baseline sleep duration for 12 hours during the day preceding deprivation and rebound sleep duration for 12 hour during the day immediately following nighttime deprivation.

### RNAi-based neurodevelopmental disorder-associated gene screen

*Drosophila* homologs of human genes of interest were identified by performing protein BLAST with the human amino acid sequences for closest *Drosophila* homologs (flybase.org/blast; annotated protein database). Homologs with alignment scores greater than 80 were included in the screen. Virgins collected from the *hs-hid; elav-GAL4; UAS-Dcr2* (*hEGD*) fly stock were crossed to males of RNAi fly stocks from the Transgenic RNAi Project (TRiP) collection(*67*). All available RNAi stocks for a given *Drosophila* gene from the TRiP collection were utilized. For controls, we used *hEGD* x TRiP library landing site host strains: P{y[+t7.7]=CaryP}attP2 (Chr3, BSC#36303) or P{y[+t7.7]=CaryP}attP40 (Chr2, BSC #36304). Female flies were loaded into the DAM system and sleep assays were performed as described. Total sleep was compared between RNAi lines and control lines to identify lines of interest.

### Circadian assays

Female flies were loaded into the DAM system 5-7 days after eclosion as described above and entrained to a 12:12: LD cycle for 3 days before being transferred to constant darkness (DD). Locomotor activity during days 2-7 in DD was analyzed in Clocklab software (Actimetrics, Wilmette, IL). Fast Fourier Transform (FFT) was performed for the locomotor activity collected during DD, and the maximum amplitude of the FFT was calculated and compared across genotypes. Flies were categorized as strongly rhythmic (FFT ≥ 0.05), moderately rhythmic (0.05 > FFT ≥ 0.03), weakly rhythmic (0.03 > FFT ≥ 0.01), or arrhythmic (< 0.01).

### Proboscis extension reflex assays

Adult female flies were collected 2-3 days post-eclosion and aged on standard fly food. 5-7 day old flies were starved for 24 hours in empty food vials on Kimwipe wet with ddH_2_O. After starvation, flies were anesthetized with CO_2_ at ZT1, and the posterior thorax and wings were gently glued to microscopy glass slides using nail polish under CO_2_ anesthesia. Flies were placed in a humidified chamber and allowed to recover for 5 hours prior to the start of the assay. For experiments, slides were mounted at a 45 degree angle under a dissecting microscope to observe proboscis extension. A 1mL syringe was used to present the following solutions to the front tarsi of each fly: ddH_2_O, 100 mM fructose, 10 mM quinine, or 1000 mM sucrose (Sigma). Flies were satiated with ddH_2_O prior to the start of the experiment. Pre-training proboscis extension was tested by presenting 100 mM fructose three times, separated by 10 second intervals. Flies that did not satiate with ddH_2_O or did not extend to initial fructose presentation (pre-training) were excluded from the remainder of the experiment. For training rounds, fructose was presented to the fly tarsi. Quinine was presented to the extended proboscis of each fly, and flies were allowed to drink for up to 2 seconds. Quinine presentation occurred in 10 second intervals within each training round. There was a one-minute interval between each training round, and a total of three training rounds before testing. For testing, fructose was presented three times to the fly tarsi with a 10 second interval between each presentation. At the end of each experiment, flies were given 1000 mM sucrose to check for intact PER, and non-responders were excluded from statistical analyses. The number of proboscis extensions was recorded during each training round and during testing, and reported as a percentage of the total number of possible extensions.

### Courtship assays

Newly-eclosed virgin male flies were collected within 4 hours after eclosion, kept in isolation on regular food, and aged to 3 days post-eclosion prior to the start of courtship experiments. Female Canton-S virgins (3-7 days post-eclosion) were used in all courtship assays. A single male and female were gently aspirated into a well-lit porcelain mating chamber (25 mm diameter and 10 mm depth) covered with a glass slide. Experiments were performed in a temperature and humidity-controlled room at 22°C, 40-50% humidity. Courtship index was determined as the percentage of time a male was engaged in courtship activity during a period of 10 minutes or until successful copulation(*68*). Courtship assays were recorded using a video camera (Sony HDR-CX405) and scored blinded to experimental condition.

### Social space assays

Newly eclosed virgin female flies were collected within 4 hours after eclosion and housed in groups of 20 in vials with standard fly food. Flies were aged to 5-7 days post-eclosion before the start of the assay. The social space arena was made of two 18 cm x 18 cm square glass plates separated by 0.47 cm acrylic spacers. Two right triangle spacers (8 cm x 16 cm) were placed on opposite sides of the square arena and two rectangular spacers (9 cm x 2 cm) were placed at the bottom of the arena, resulting in an isosceles triangle-shaped space (base: 15.2 cm and height: 15.2 cm). Flies were gently aspirated into the social space arena by briefly removing the bottom rectangular spacers. Forty flies were included in each assay. After all 40 flies were introduced to the arena, the rectangular spacers were replaced, and the bottom of the arena was firmly tapped down 5 times. Digital images were captured using a video camera (Sony HDR-CX405) after allowing flies to settle for 20 minutes. Images were imported into Fiji for analysis. For each image, a body length measurement was taken as the average length in pixels, measured from top of the head to tip of the abdomen, of 5 randomly selected flies within the arena. The center of each fly was manually selected, and an automated measure of the nearest neighbor to each selection was determined using the Nearest Neighbor Distances Calculation plugin on Fiji (https://icme.hpc.msstate.edu/mediawiki/index.php/Nearest_Neighbor_Distances_Calculation_with_ImageJ). Results were binned by body length distances. We calculated a social space index (SSI) as the difference between the number of flies in the first bin (0-2 body lengths) and the number of flies in the second bin (2-4 body lengths).

### TARGET system experiments

For development temporal mapping experiments using the TARGET system, parental crosses were maintained on standard fly food at 18°C. Timed egg lays were achieved by flipping parental crosses to bottles with standard fly food from ZT1-ZT6 at 28°C, or from ZT1-ZT8 at 18°C. To achieve *ISWI* knockdown at certain stages, flies were kept at 28°C to allow for GAL80 denaturation. To repress RNAi expression, flies were moved to 18°C to prevent GAL80 denaturation. Due to temperature-related changes in *Drosophila* developmental timing, developmental periods were visually determined. Genetic controls were subject to the same temperature shifts as experimental flies to account for the effect of changing temperature on development. Sleep assays were conducted on 5-7 day old female flies at 22°C in 12:12 LD.

### 2^nd^ instar larval sleep experiments

To synchronize developmental stages, adult fly parental crosses were placed in embryo collection cages (Genesee Scientific, cat#: 59-100) for 24 hours. Eggs were laid on a petri dish containing 3% agar, 2% sucrose, and 2.5% apple juice with yeast paste spread on top. Molting 1^st^ instar larvae were collected two days after egg lay and moved to a separate petri dish with yeast and allowed to molt into 2^nd^ instar. Freshly molted 2^nd^ instar larvae were placed in the LarvaLodge to monitor sleep as previously described. Collected data were analyzed using a custom MATLAB code(*44*).

### Thermogenetic activation experiments

Animals were reared at 18°C to prevent activation of TrpA1 during development. Adult female flies were collected 2-3 days after eclosion, and aged at 18°C on standard fly food. 5-7 day old flies were loaded into the DAM system to monitor sleep and placed at 22°C on a 12:12 LD schedule for 3 days. TrpA1 activation was achieved by a temperature shift to 31°C across non-consecutive 12-hour light or 12-hour dark period. Between increases in temperature, flies were kept at 22°C.

### RNA-Seq

#### Dissection and RNA extraction

40 brains per sample at the mid-3^rd^ instar stage were dissected in cold AHL (108 mM NaCl, 5 mM KCl, 2 mM CaCl_2_, 8.2 mM MgCl_2_, 4mM NaHCO_3_, 1 mM NaH_2_PO_4_-H_2_O, 5 mM trehalose, 10 mM sucrose, 5 mM HEPES). Three biological replicates for the control group and four for the experimental, each with 40 brains, were dissected. Brains were transferred to 1 mL of Trizol and incubated for 5 minutes at room temperature (RT). 0.2 mL of chloroform was added and samples were inverted. Samples were incubated 2-3 minutes at RT, then centrifuged at 12,000g for 15 minutes at 4°C. Genomic DNA was removed using a gDNA eliminator column (RNeasy Plus Micro Kit, Qiagen). RNA was then extracted using the RNeasy MinElute Cleanup Kit (Qiagen).

#### RNA library preparation and sequencing

Sequence libraries for each sample were synthesized using the NEBNext Ultra II Directional RNA kit following supplier recommendations and were sequenced on Illumina HiSeq-4000 sequencer as single reads of 100 base reads following Illumina’s instructions.

#### Differential gene expression analysis

RNA-seq reads were mapped to the *Drosophila melanogaster* assembly BDGP6 pre-indexed with transcript models from Ensembl 87 using STAR 2.5.0b with default parameters except – alignIntronMax set to 10,000. Aligned reads were assigned to gene models using the summarizeOverlaps function of the GenomicRanges R package. Reads per kilobase per million (RPKMs) were calculated with a slight modification, whereby only reads assigned to annotated protein-coding genes were used in the denominator, to minimize batch variability due to different amounts of contaminant ribosomal RNA. Differential expression was determined using the DESeq2 package. The annotated genes exhibiting an adjusted P-value > 0.1 were considered to be differentially expressed compared to control. Visualization of differentially expressed genes was done using R-package ggplot2 v3.2.0.

#### Gene interaction network analysis

Human homologs of *Drosophila* DEG in the setting of *ISWI* knockdown were identified using DIOPT (DRSC Integrative Ortholog Prediction Tool) v.8.0(*69*). We assessed the connectivity of *SMARCA5* and *SMARCA1* with these DEG homologs in the context of a brain-specific gene interaction network(*56, 57*). This network was constructed using a Bayesian classifier trained on gene co-expression data, which predicts the likelihood of interactions between pairs of genes in the brain. We generated a sub-network containing all interactions with weights >2.0 (the top 0.5% of all pairwise interactions) from the entire interaction network. Using the NetworkX Python package, we next identified the shortest distances as a measure of connectivity, as well as connector genes within the shortest paths, between the *SMARCA* genes and DEGs(*70*). We similarly assessed the connectivity of *SMARCA* genes with 224 human genes annotated for sleep and circadian rhythm functions (Gene Ontology terms GO:0007623 and GO:0030431). Network diagrams were generated using Cytoscape v.3.7.2(*71*)^71^, and Gene Ontology enrichment analysis of the connector genes was performed using the PantherDB Gene List Analysis tool(*72*).

#### Immunohistochemistry

Fly brains were dissected in 1xPBS with 0.1% Triton-X 100 (PBST) and fixed in 4% PFA for 15 minutes at room temperature. Following 3×10 minute washes in PBST, brains were incubated with primary antibody at 4°C overnight. Brains were washed 3×10 minutes in PBST, and incubated with secondary antibody for 2 hours at room temperature. After 3×10 minute PBST washes, brains were cleared in 50% glycerol and mounted in Vectashield. The following primary antibodies were used at 1:1000 dilutions: mouse 1D4 anti-Fasciclin II (anti-FasII; Developmental Studies Hybridoma Bank), rabbit anti-HA-Tag (Cell Signaling), guinea pig anti-PER (a gift from Dr. Amita Seghal) and rabbit anti-GFP (Thermo Fisher). The following secondary antibodies were used at 1:1000 dilutions: Alexa Fluor 488 Donkey anti-mouse, Alexa Fluor 488 Donkey anti-rabbit, Alexa Fluor 488 Donkey anti-guinea pig, and Alexa Fluor 647 Donkey anti-mouse (Thermo Fisher).

#### Imaging and analysis

Microscopy images were taken using a Leica TCS SP8 confocal microscope. Images were processed in NIH Fiji. All settings were kept constant between conditions within a given experiment. Images were taken in 1.0um steps unless otherwise noted.

1. <u>PER quantification</u> To investigate PER expression in sLNVs and lLNVs, brains were co-stained with anti-PDF (to label relevant cells) and anti-PER antibodies. Brains were dissected at CT0, CT4, CT12, and CT20. We defined the area of each sLNV or lLNV cell body by PDF staining. Area, mean gray value, and integrated density of the PER signal was measured for each cell body. Corrected total cell fluorescence (CTCF) of the cell body was calculated with the formula: CTCF = Integrated density_cell_ – (Area_cell_ x Mean background fluorescence). All cells per brain were averaged and compared across genotypes.
2. <u>MB quantification</u> For temporal mapping, spatial mapping (*OK107-GAL4 > UAS-ISWI RNAi*), and neuroblast experiments, a maximum projection image of all Z-slices was generated. MB morphology was manually quantified from the maximum projection as a binary normal vs abnormal based on anti-FasII staining and prior description of normal MB morphology. For *SMARCA1^WT^* and *SMARCA5^WT^* rescue experiments, for each Z-slice, the vertical or horizontal lobe on one hemisphere was manually outlined. The full volume of the vertical or horizontal lobe was measured using the 3D Objects counter function in Fiji with the following settings: threshold = 1 and minimum puncta size = 100.
3. <u>dFB volume</u> For each Z-slice, the dFB was selected based on anti-GFP staining for *R23E10* dFB projections. The full volume of the dFB was measured using the 3D Objects Counter function in Fiji with the following settings: threshold = 1 and minimum puncta size = 10000.
4. <u>dFB cell body count</u> GFP-positive soma were counted across the entire brain based on anti-GFP staining for *R23E10* neurons.

#### Statistical Analysis

All statistical analyses were performed using GraphPad Prism (version 8.4.1). Sample size, specific tests, and significance values are denoted in figure legends.

## Acknowledgements

We thank members of the Kayser lab, Drs. Maja Bucan, Roberto Bonasio, Thomas Jongens, David Raizen, and members of the Penn Chronobiology and Sleep Institute for helpful discussions/input, and the Next Generation Sequencing Core (University of Pennsylvania) for sequencing/analysis support. We thank Dr. Roberto Bonasio for bioinformatic support in analyzing RNA Seq data. Figure S9a was created with BioRender.com. We thank Salina Yuan for assistance with creating Figure S9b.

## Author contributions

N.N.G., L.C.D, and M.S.K. conceived the project. N.N.G., L.C.D., E.H.M., and M.S.K. designed experiments. N.N.G., L.C.D., C.E.W., and M.S. performed experiments and interpreted data. M.J. and S.G. performed bioinformatic network analysis experiments. T.Y.T., M.A.D., and D.L. identified patient *SMARCA5* mutations. N.N.G., L.C.D., Q.W., and Y.S. generated *Drosophila* reagents. N.N.G. and M.S.K. wrote the manuscript, with input from all authors.

## Data and materials availability

Additional data related to the paper may be requested from the authors.

## Supplementary Materials

**Fig. S1:**
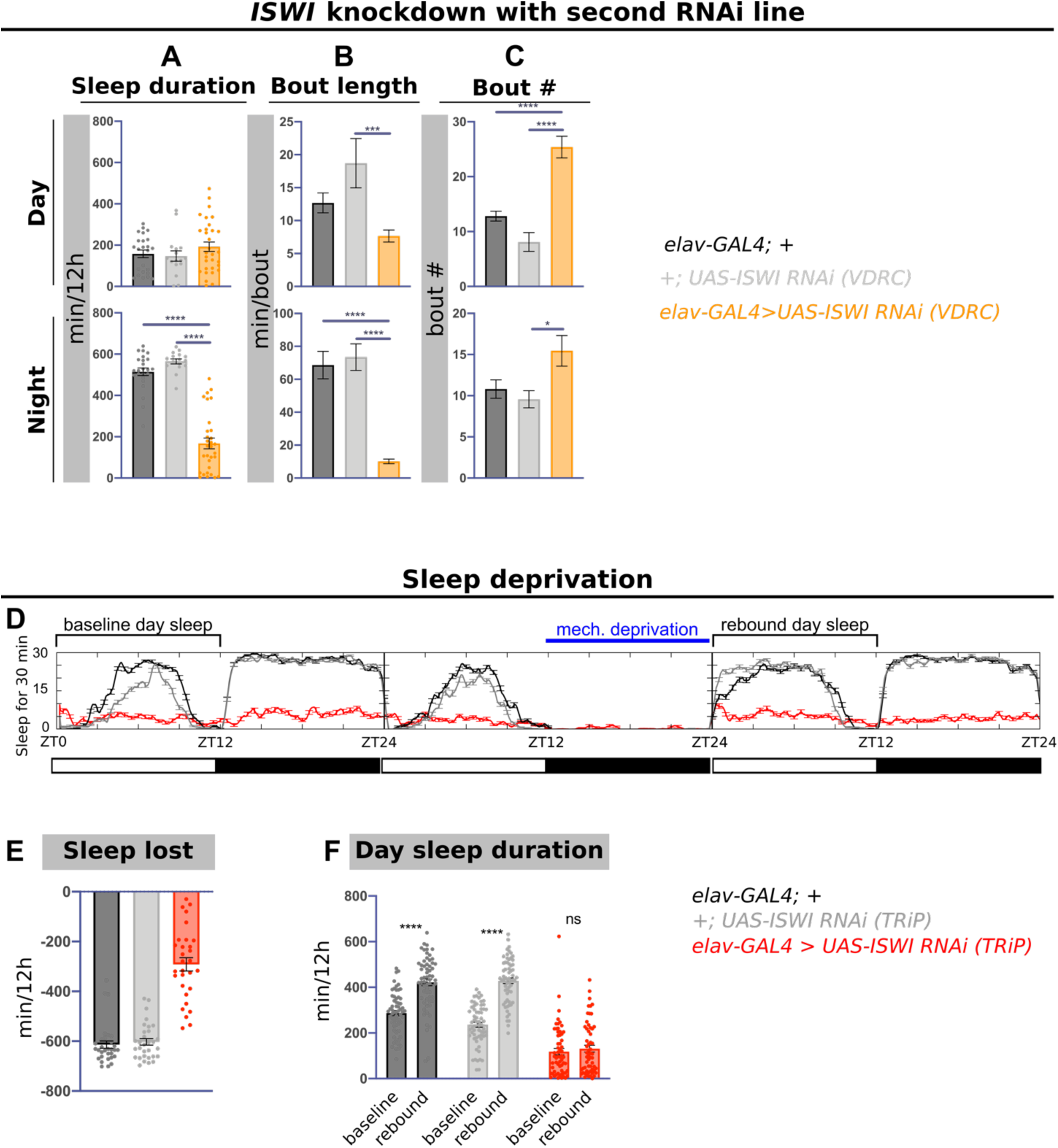
Extended characterization of sleep deficits in the setting of *ISWI* knockdown. Day and night average A) sleep duration, B) bout length, and C) bout number the VDRC *ISWI* RNAi line (orange) compared to genetic controls (black and gray) (n = 24, 16, 31 from left to right). D) Representative sleep traces for mechanical sleep deprivation experiment. E) Nighttime sleep loss in minutes during mechanical deprivation compared to baseline nighttime sleep in *ISWI* knockdown (red) and genetic controls (black and gray). F) Comparison of baseline day sleep before night time mechanical deprivation and rebound day sleep directly following deprivation (n = 63, 65, 61 from left to right; mixed model ANOVA with post-hoc Tukey’s multiple comparison test).

**Fig. S2:**
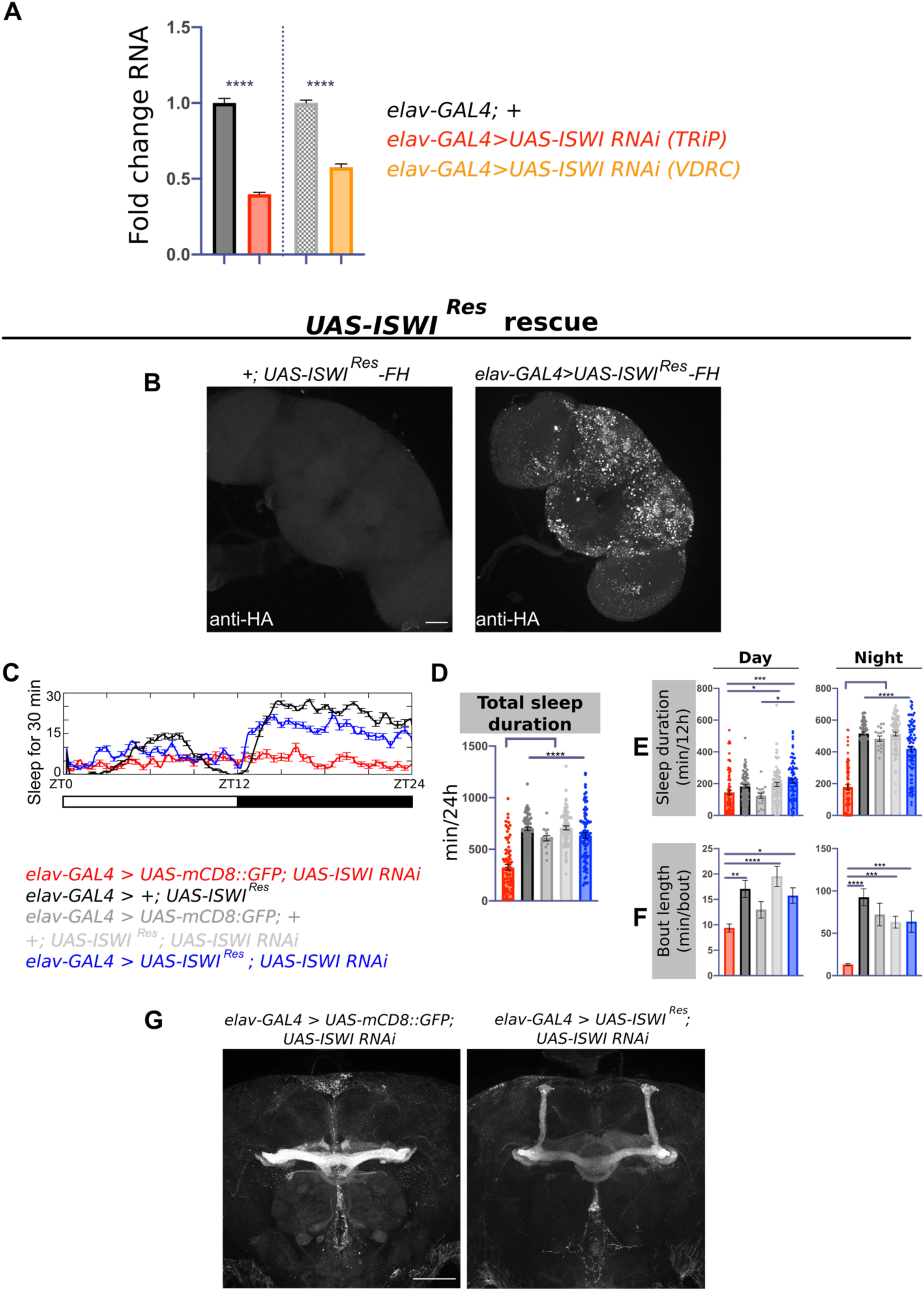
Validation of *ISWI* knockdown. A) qPCR for *ISWI* mRNA in the setting of knockdown (n = 3 replicates, ≥10 brains per genotype per replicate). B) Immunostaining for HA in adult fly brains confirms FLAG and HA-tagged *UAS-ISWI^Res^* is expressed. Negative control (left) compared to *elav-GAL4>UAS-ISWI^Res^-FLAG-HA.* Scale bar, 50 um. C) Representative sleep traces of *ISWI* knockdown (red), *ISWI^Res^* overexpression (black), and *ISWI^Res^* rescue (blue). D) Total sleep in *ISWI^Res^* rescue compared to controls (n = 77, 71, 16, 81, 88 from left to right). Day and night average E) sleep duration and F) sleep bout lengths in *ISWI^Res^* rescue compared to controls. G) Example FasII immunostaining of adult fly mushroom bodies from *ISWI* knockdown (left) and *ISWI^Res^* rescue (right). Scale bar, 50 um.

**Fig. S3:**
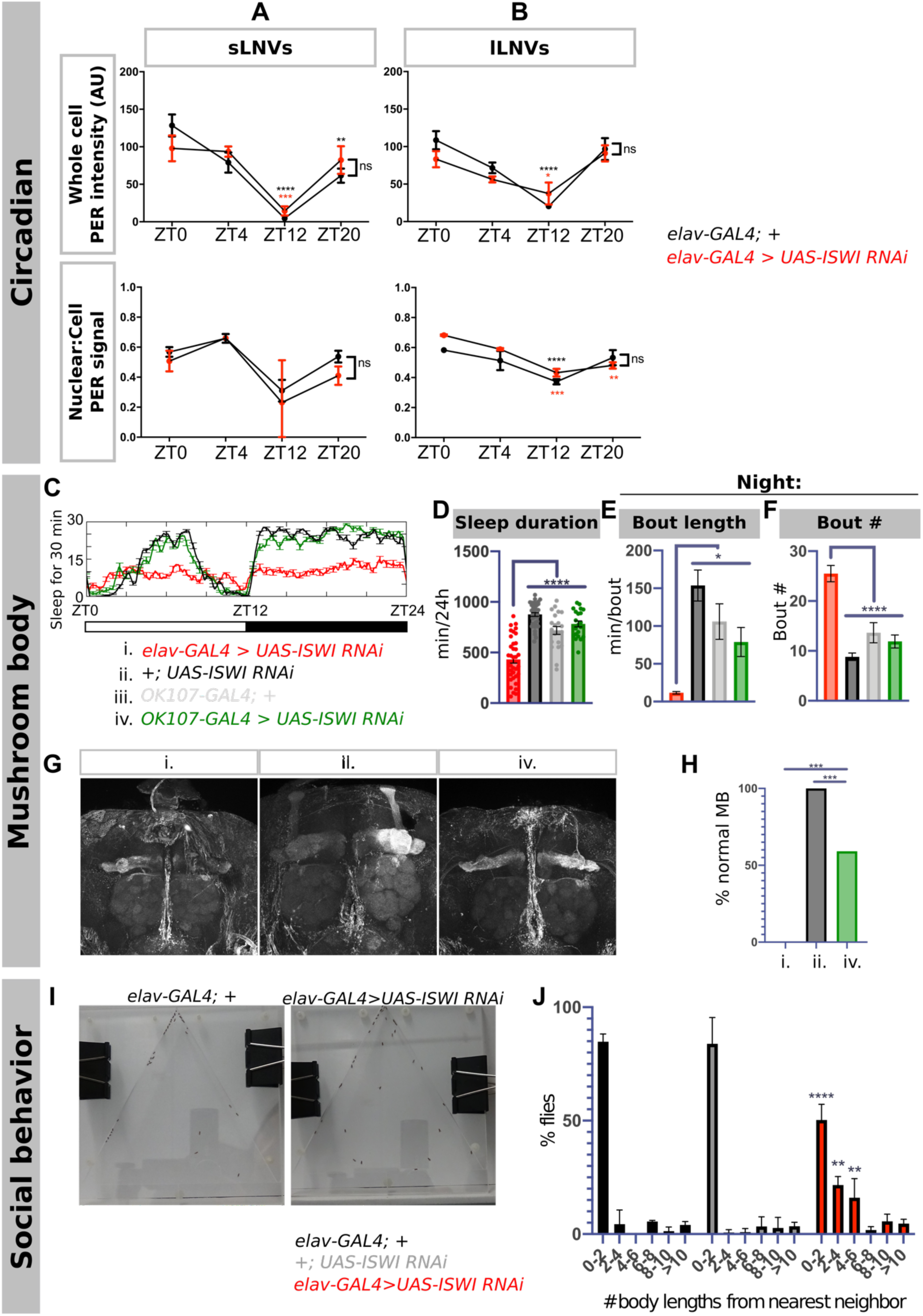
Extended characterization of circadian, mushroom body morphology, and social behavior deficits in the setting of *ISWI* knockdown. Quantification of PER staining intensities at ZT0, ZT4, ZT12, and ZT20 in A) small LNVs (sLNVs) and B) large LNVs (lLNVs). Whole cell PER staining intensity (top graphs) vs nuclear:cell PER signal (bottom graphs) (n ≥ 5 brains per genotype per timepoint) (two-way ANOVA; post-hoc Bonferroni’s multiple comparison tests. Asterisks represents significant differences compared to ZT0 for each genotype; brackets represent comparison between groups, no significant difference noted at each timepoint). C) Example sleep traces comparing *OK107-GAL4 > UAS-ISWI RNAi* (green), pan-neuronal *ISWI* knockdown (red), and genetic controls (black and gray). D) Total sleep duration and nighttime E) sleep bout length and F) sleep bout number in *OK107-GAL4 > UAS-ISWI RNAi* compared to genetic controls and *elav-GAL4 > UAS-ISWI RNAi* (n = 36, 40, 21, 21 from left to right). Significance bars across multiple groups represent significant comparisons across all groups compared to pan-neuronal knockdown. G) Example FasII immunostaining of adult fly brains in the setting *ISWI* knockdown using *elav-GAL4* (left), genetic control (middle), and *OK107-GAL4* (right). H) Quantification of FasII staining in *OK107-GAL4 > UAS-ISWI RNAi* as a binary of normal or abnormal MB morphology (n = 20, 14, 22 from left to right) (pairwise Fisher’s Exact Test). I) Sample image of social space arenas for each condition. J) Histogram distribution of body length distances in social space arenas (n ≥ 3 replicates per genotype, 40 flies per replicate per genotype) (Two-way ANOVA with post-hoc Tukey’s multiple comparisons test; asterisks denote significant bins in the *elav-GAL4 > UAS-ISWI RNAi* group compared to both genetic controls in post-hoc testing).

**Fig. S4:**
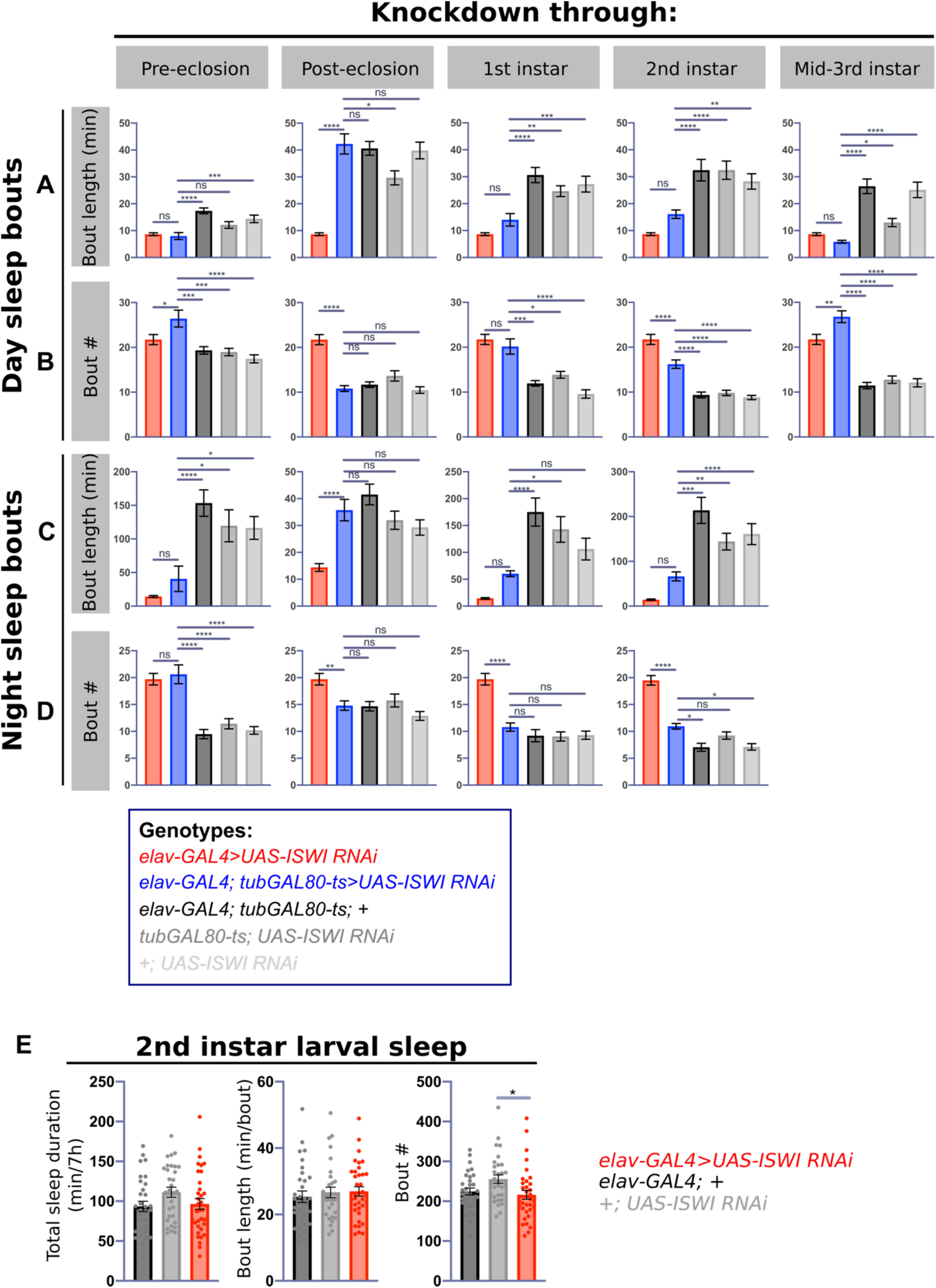
Extended sleep characterization in temporally restricted *ISWI* knockdown and during larval development. Quantification of average daytime A) bout length and B) bout number, and average nighttime C) bout length and D) bout number for *ISWI* knockdown through pre-eclosion, post-eclosion, 1^st^ instar, 2^nd^ instar, and mid-3^rd^ instar (left to right) (sample sizes for each knockdown condition are denoted in the legend for figure 3). E) Quantification of total sleep during a 7-hour period during 2^nd^ instar in the setting of *ISWI* knockdown (n = 31, 31, 55 from left to right).

**Fig. S5:**
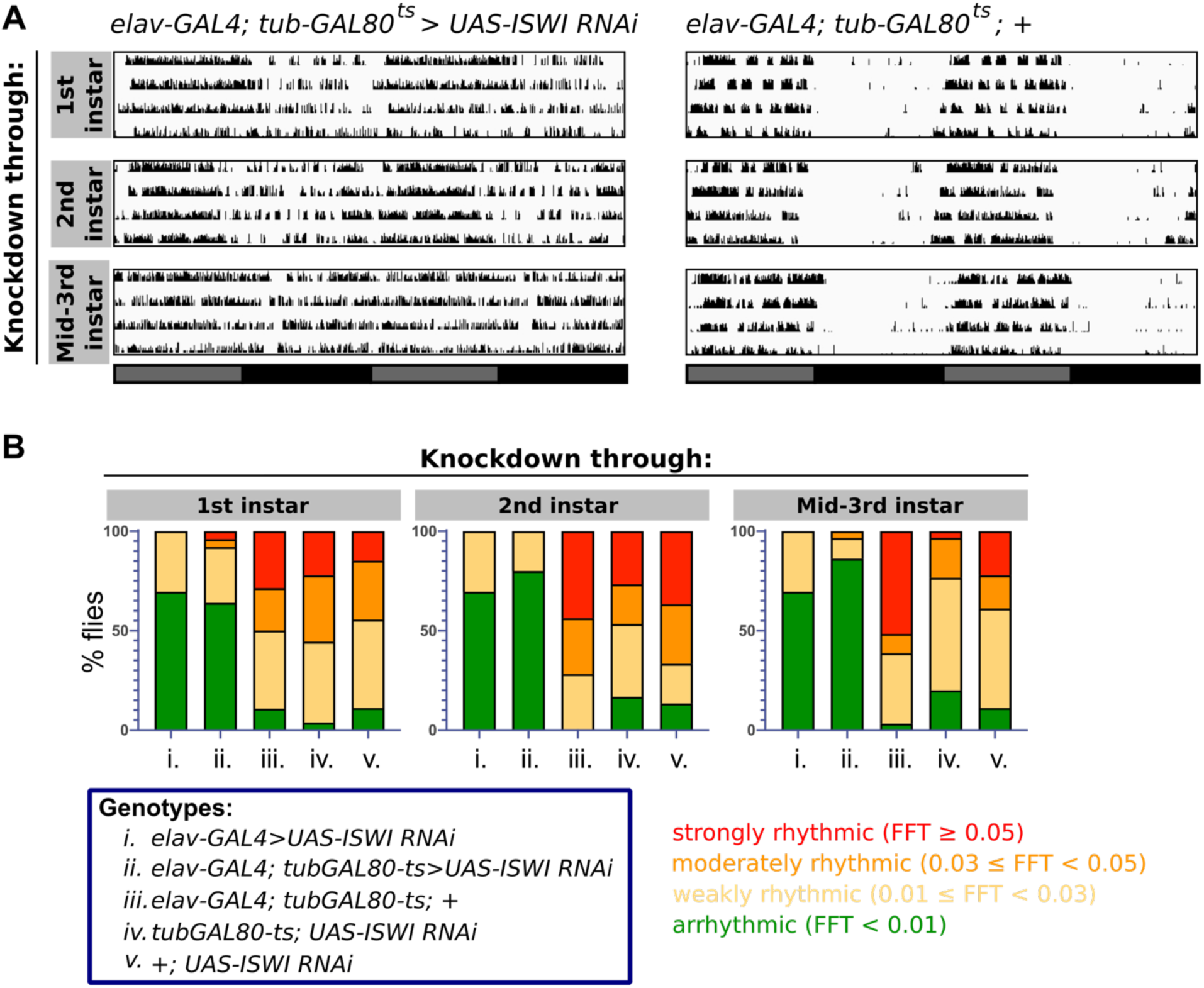
Extended characterization of circadian rhythms in the setting of temporally restricted *ISWI* knockdown. A) Representative actograms of temporally-restricted *ISWI* knockdown (left) through 1^st^, 2^nd^, or mid-3^rd^ instar (top to bottom), compared to a genetic control exposed to the same temperature shifts (right). B) Proportion of strongly rhythmic, moderately rhythmic, weakly rhythmic, and arrhythmic flies in the setting of *ISWI* knockdown at different points in development compared to constitutive knockdown and in genetic controls.

**Fig. S6:**
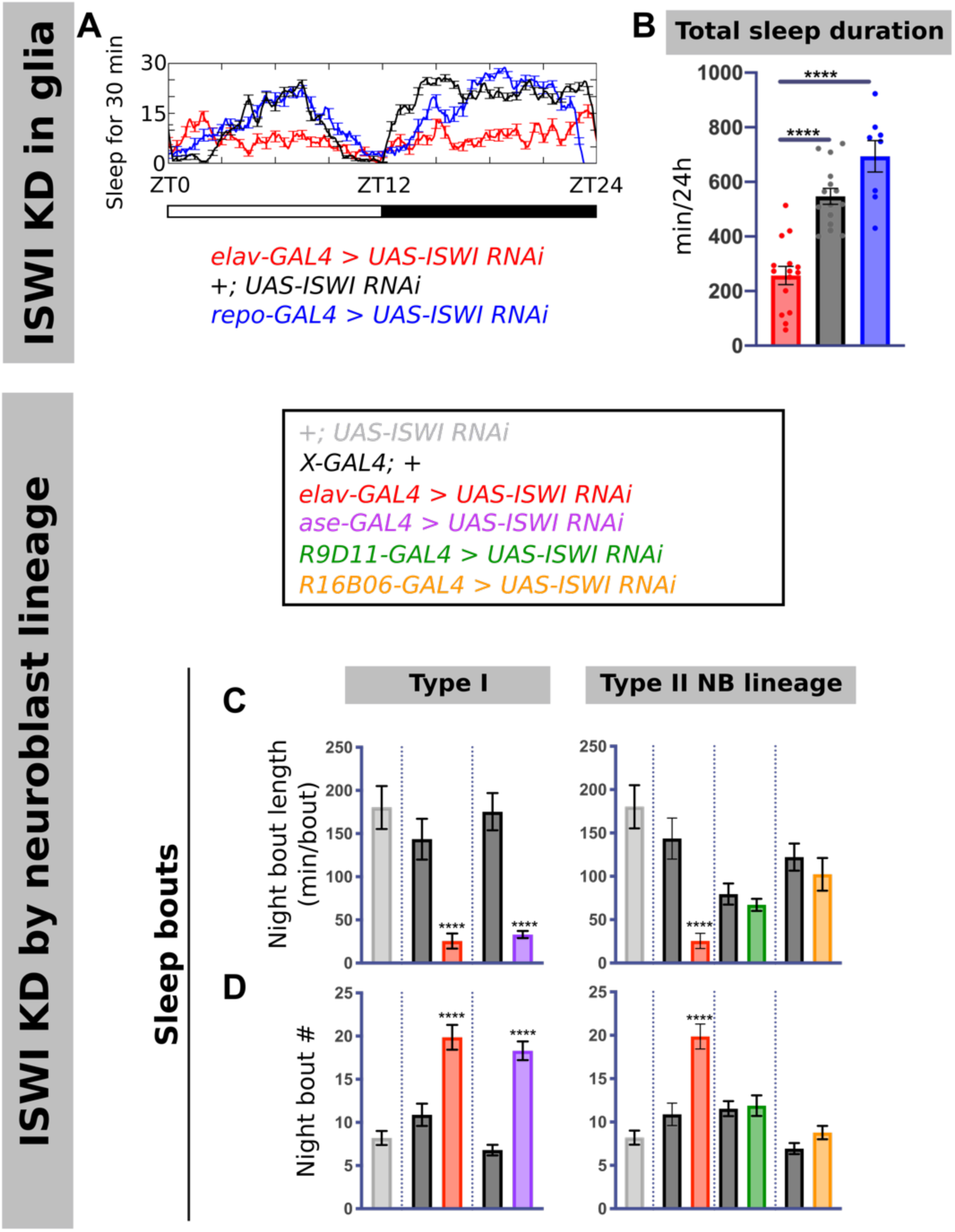
*ISWI* knockdown in glia and extended sleep characterization of knockdown in neuroblast lineages. A) Representative sleep traces and B) quantification of total sleep duration with *ISWI* knockdown in glia (blue) compared to genetic control (black) and *elav-GAL4* knockdown (red) (n = 15, 15, 8 from left to right). Nighttime C) sleep bout length and D) sleep bout number with *ISWI* knockdown in different neuroblast lineages (for sample sizes see figure 5 legend).

**Fig. S7:**
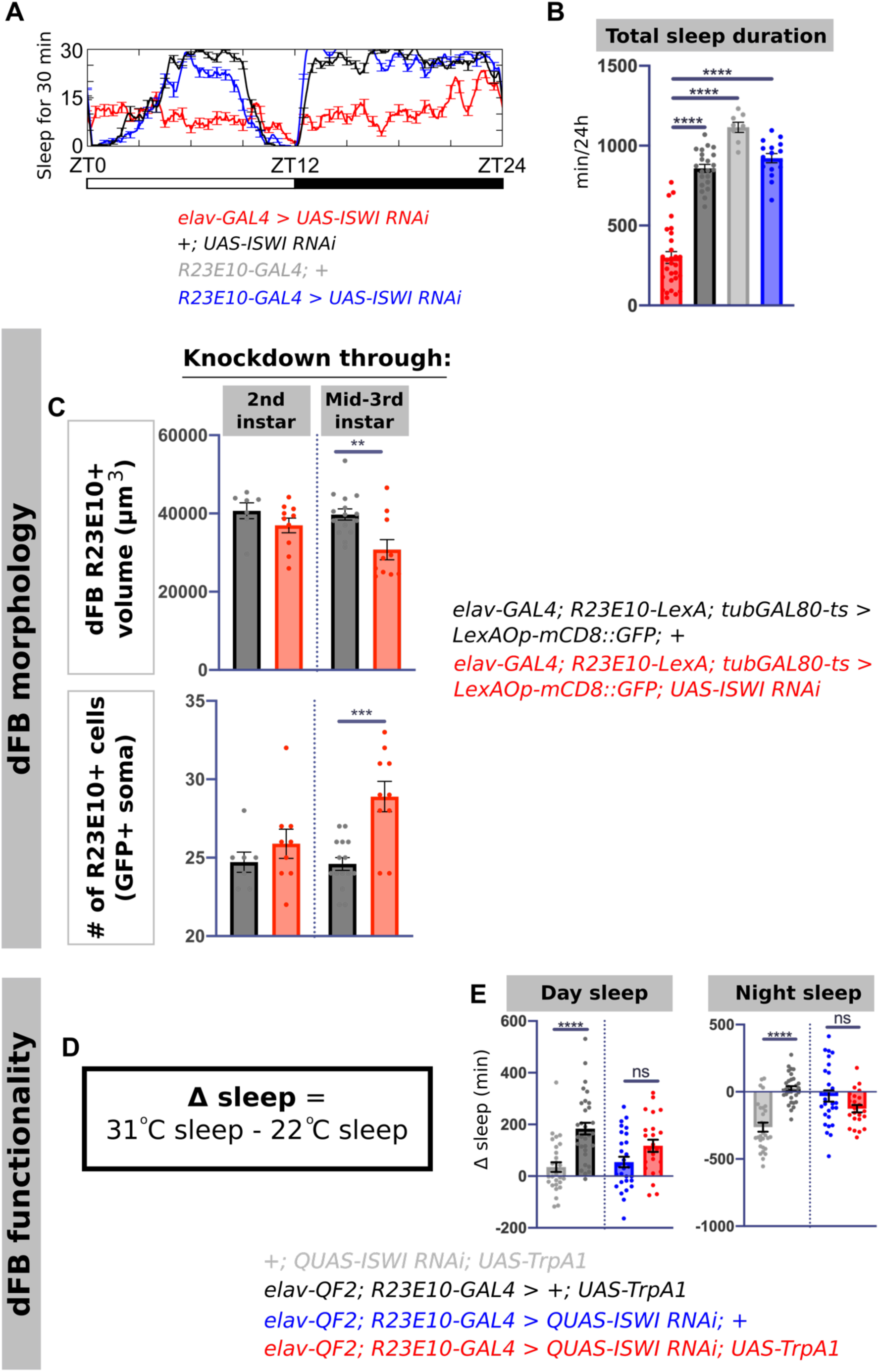
Further characterization of *R23E10* neurons in the setting of *ISWI* knockdown. A) Representative sleep traces and B) quantification of total sleep in *R23E10-GAL4 > UAS-ISWI RNAi* (blue) compared to genetic controls (black and gray) and pan-neuronal *ISWI* knockdown (red) (n = 29, 22, 8, 16 from left to right). C) Quantification of dFB volume (top) and number of *R23E10* soma (bottom) as measured by GFP immunostaining in temporally-restricted *ISWI* knockdown (red) compared to genetic controls exposed to the same temperature shifts (black) (n = 7, 10, 15, 10 brains from left to right). D) Formula to calculate sleep differences per individual fly resulting from R23E10 thermogenetic activation. E) Day (left) and night (right) individual fly sleep duration differences resulting from thermogenetic activation of R23E10 neurons, in the setting of *ISWI* knockdown (red) compared to negative controls (lacking *23E10>TrpA1*) with and without pan-neuronal *ISWI* knockdown (gray and blue) and positive control (with *23E10>TrpA1*) without *ISWI* knockdown (black) (see figure 6 legend for sample size).

**Fig. S8:**
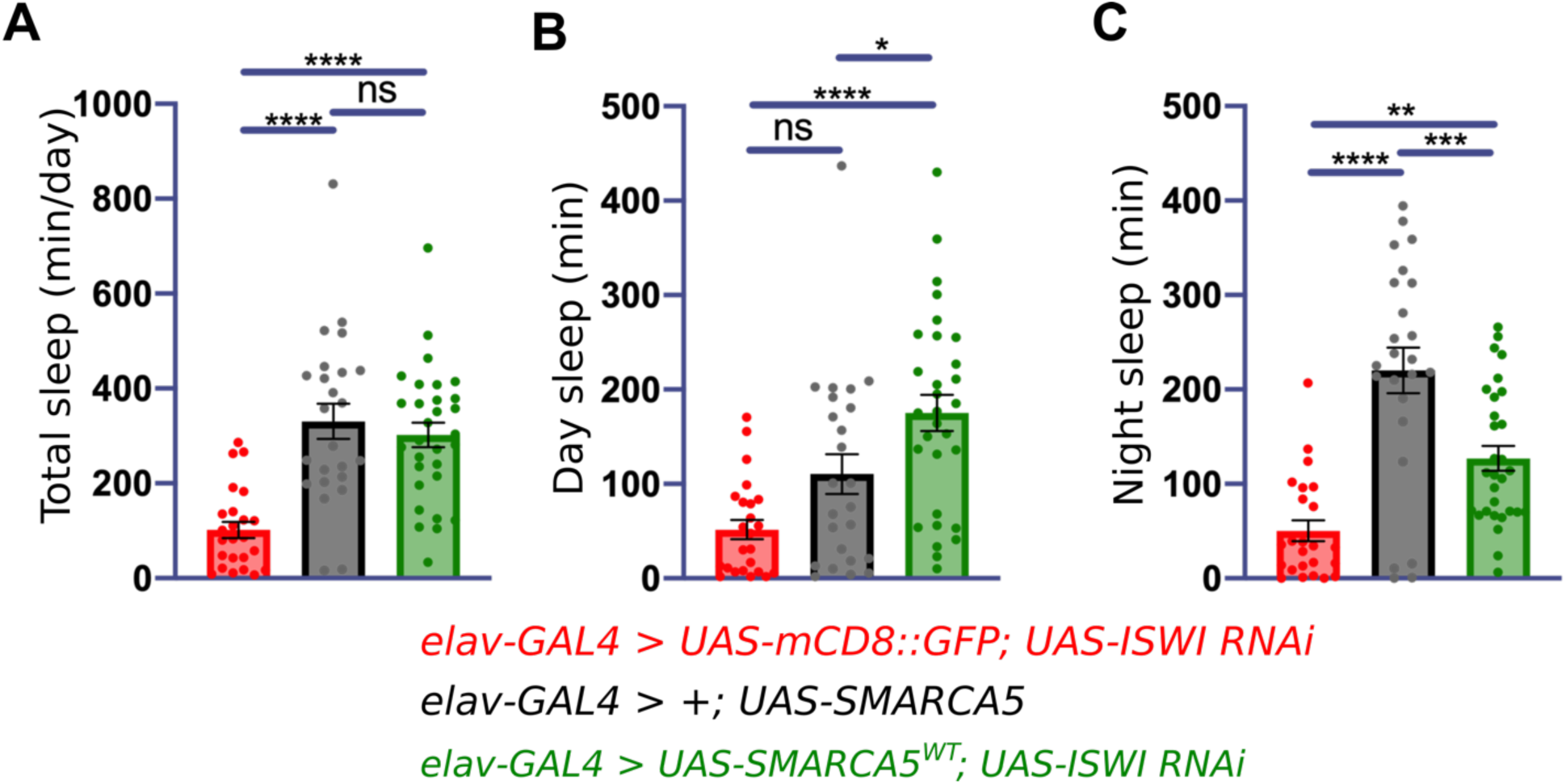
High resolution sleep monitoring for *SMARCA5^WT^* rescue in the setting of *ISWI* knockdown confirms sleep rescue. Quantification of A) total sleep duration, B) day sleep duration, and C) night sleep duration from multibeam monitoring of *SMARCA5^WT^* rescue (green) in the setting of pan-neuronal *ISWI* knockdown, along with pan-neuronal *ISWI* knockdown (red), and *SMARCA5^WT^* overexpression alone (black) (n = 24, 24, 30 from left to right).

**Fig. S9:**
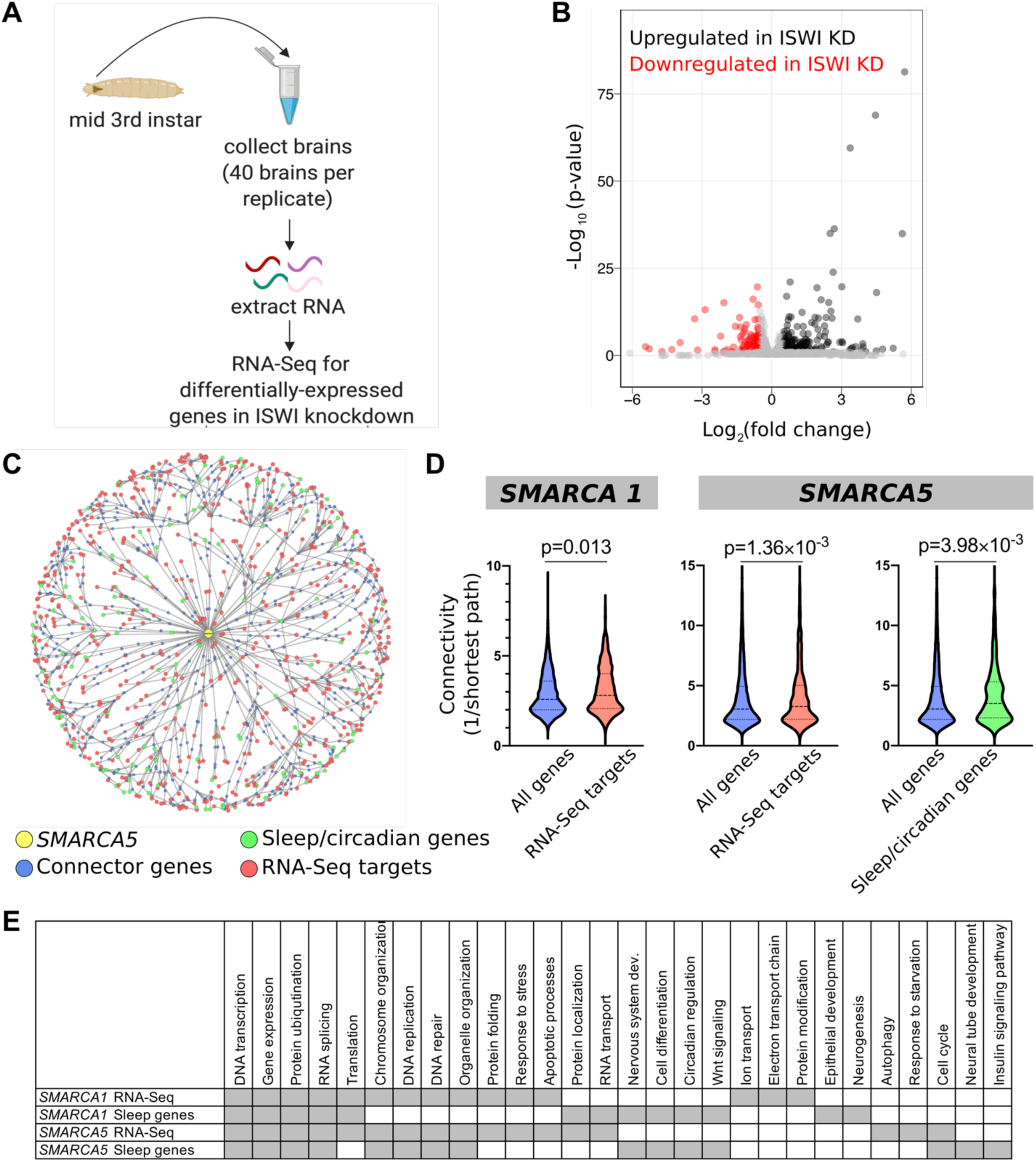
Gene network analysis reveals increased connectivity between *SMARCA5* and human genes regulating sleep and circadian functions. A) RNA-Seq experiment design. B) Visualization of differentially expressed genes (upregulated – gray, downregulated – red) in the setting of *ISWI* knockdown (n ≥ 3 replicates per genotype, ≥ 40 larval brains per replicate; see methods for statistical tests, adjusted P > 0.1). *ISWI* (P-adj = 5.14 x 10^−230^) and *dsx* (P-adj = 1.27 x 10^−109^) were not included in the graph to facilitate visualizing other genes. C) Visualized connectivity between *SMARCA5* (yellow), human orthologs of *Drosophila* RNA-Seq targets (red), and human sleep and circadian genes (blue). Connector genes are denoted in green. D) Quantification of connectivity (inverse of shortest path length) between *SMARCA1* and *SMARCA5* and all genes in the network (blue) and RNA-Seq targets (red), as well as between *SMARCA5* and human sleep and circadian genes (green) (two-tailed T-tests). E) Representative enriched GO Biological Process terms (P<0.05, Fisher’s Exact test with Benjamini-Hochberg correction) among connector genes between *SMARCA1* and *SMARCA5* and RNA-Seq target genes or human sleep and circadian genes.

**Table S1:**
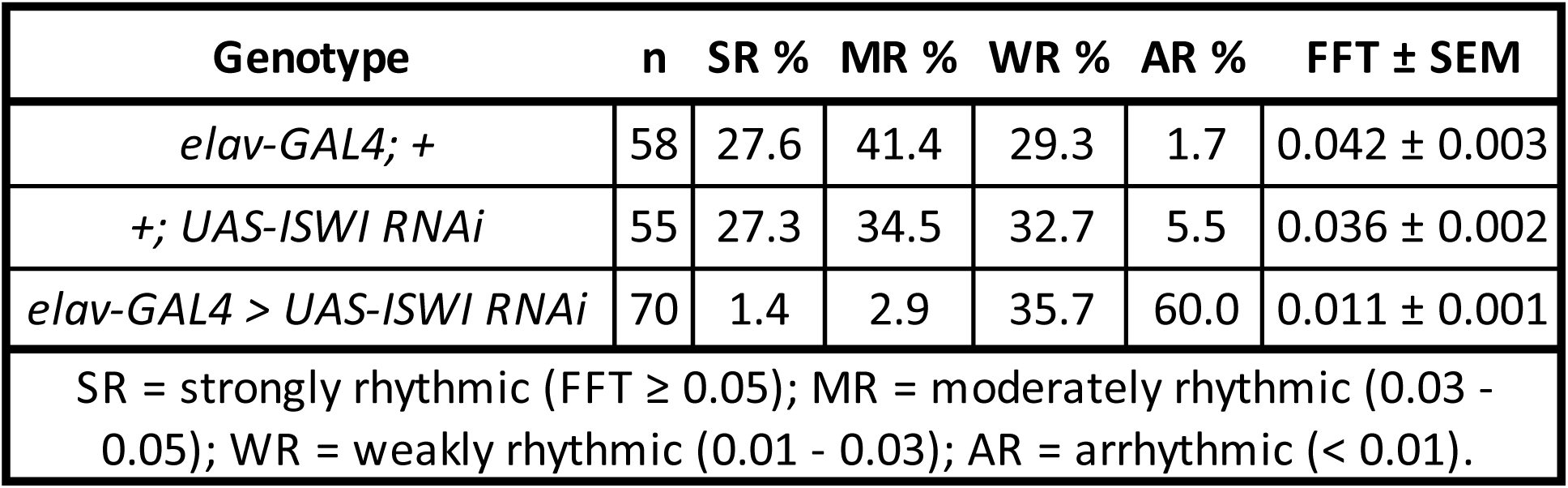
Percentage of strongly rhythmic, moderately rhythmic, weakly rhythmic, and arrhythmic flies in *elav-GAL4 > UAS-ISWI RNAi* flies and in genetic controls as defined by FFT amplitude. Mean FFT amplitude with SEM shown for each genotype.

**Table S2:**
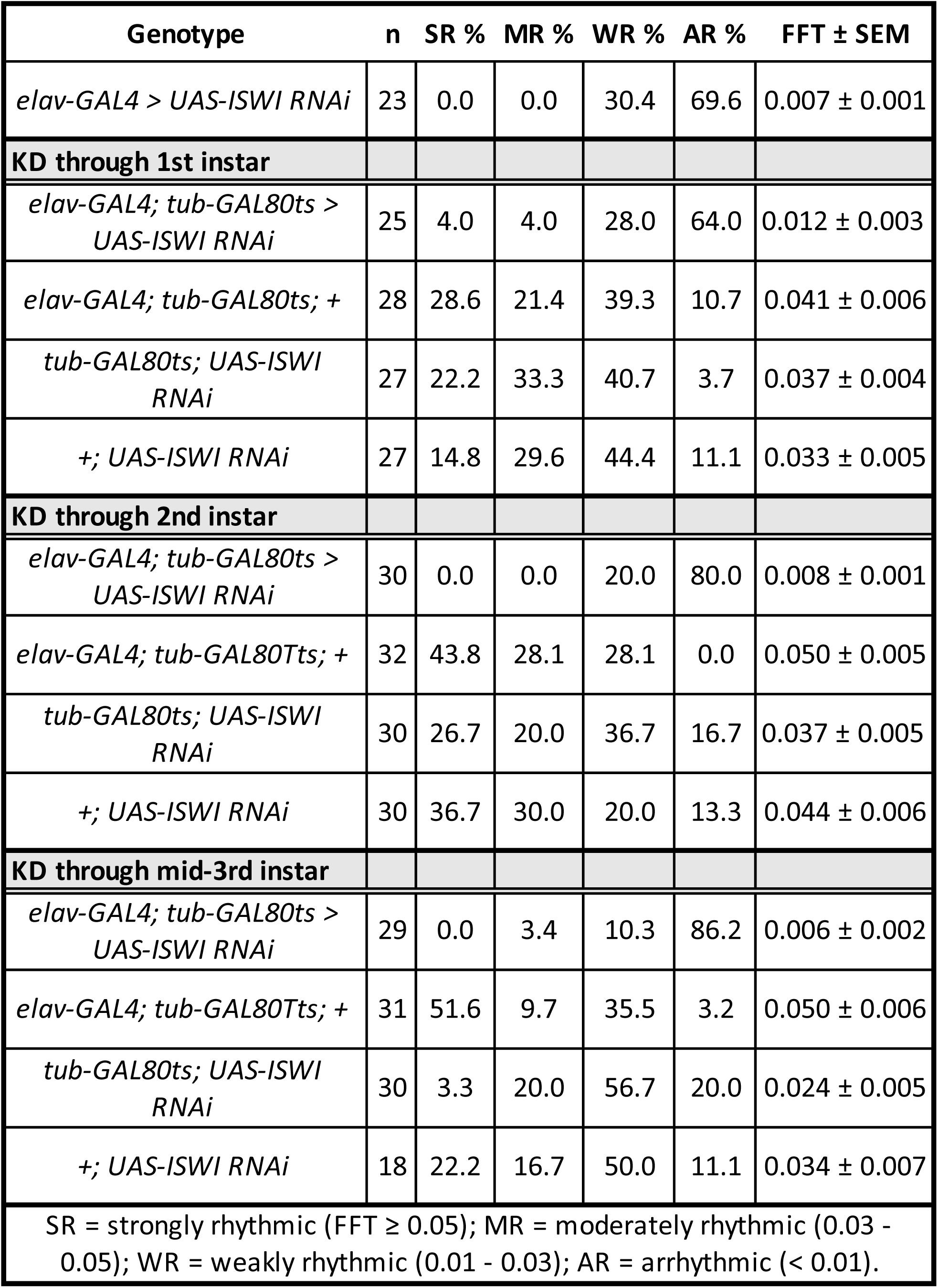
Percentage of strongly strongly rhythmic, moderately rhythmic, weakly rhythmic, and arrhythmic flies as defined by FFT amplitude for temporally restricted *ISWI* knockdown through the noted time points. Mean FFT amplitude with SEM shown for each genotype and temporal knockdown condition.

